# Cross-species single-cell comparison of systemic and cardiac inflammatory responses after cardiac injury

**DOI:** 10.1101/2023.03.15.532865

**Authors:** Eric Cortada, Jun Yao, Yu Xia, Friederike Dündar, Paul Zumbo, Boris Yang, Alfonso Rubio-Navarro, Björn Perder, Miaoyan Qiu, Anthony M. Pettinato, Edwin A. Homan, Lisa Stoll, Doron Betel, Jingli Cao, James C. Lo

## Abstract

The immune system coordinates the response to cardiac injury and is known to control regenerative and fibrotic scar outcomes in the heart and subsequent chronic low-grade inflammation associated with heart failure. Here we profiled the inflammatory response to heart injury using single cell transcriptomics to compare and contrast two experimental models with disparate outcomes. We used adult mice, which like humans lack the ability to fully recover and zebrafish which spontaneously regenerate after heart injury. The extracardiac reaction to cardiomyocyte necrosis was also interrogated to assess the specific peripheral tissue and immune cell reaction to chronic stress. Cardiac macrophages are known to play a critical role in determining tissue homeostasis by healing versus scarring. We identified distinct transcriptional clusters of monocytes/macrophages in each species and found analogous pairs in zebrafish and mice. However, the reaction to myocardial injury was largely disparate between mice and zebrafish. The dichotomous response to heart damage between the mammalian and zebrafish monocytes/macrophages may underlie the impaired regenerative process in mice, representing a future therapeutic target.

## Introduction

Cardiovascular disease is the leading cause of death worldwide with myocardial infarction (MI) and its complications including congestive heart failure (CHF) accounting for the lion’s share of the burden^1, 2^. Despite the local nature of the injury, the adverse effects of MI are not restricted to the cardiovascular system. Survivors of MI often experience liver and kidney injury, fever, inflammation and have increased risk of certain cancers and ischemic stroke^3^ ^4, 5^. In humans and other mammals, MI resulting from an acute disruption in coronary artery blood flow to the heart leads to cardiomyocyte necrosis. Due to the incapacity of the adult mammalian heart to regenerate, the heart is ultimately repaired through a predominantly fibrotic process with many patients suffering from CHF.

It has long been known that MI and heart failure with reduced ejection fraction (HFrEF) are associated with chronic low-grade inflammation^6^. That chronic inflammation after MI can drive adverse cardiovascular events has been elegantly shown in clinical trials with blockade of some inflammatory pathways^7, 8^. Post-MI inflammation may also contribute to the pathogenesis of breast and lung cancer, and chronic liver and kidney disease^9–12^. The dynamic inflammatory process post MI evolves from an acute phase characterized by the elaboration of proinflammatory cytokines and chemokines, immune cell infiltration to the site of injury, and systemic leukocytosis to a resolution or reparative phase punctuated by a tissue clearing and fibrotic cell and gene program in the heart with unresolved systemic inflammation in some patients.

Unlike mammals, adult zebrafish possess a remarkable capacity for cardiac regeneration with minimal scarring. This is achieved through proliferation of spared cardiomyocytes with cellular and molecular supports from non-muscle cells including immune cells^13, 14^. Cryoinjury of the zebrafish heart triggers a sequence of dynamic pro- and anti-inflammatory programs that govern successful myocardial regeneration. Swift neutrophil infiltration to the injury site in the acute inflammation stage (within 1 day of injury) is followed by macrophage recruitment that spans the inflammatory and regenerative stages (until 7 days after injury), while NK or T cells take on activity during the regeneration stage^15, 16^. This process employs various, functionally diverse populations of macrophages that play vital roles for zebrafish heart regeneration^16–18^. Early macrophage infiltration of the wound is critical for successful regeneration and results in transient fibrosis through direct and indirect contributions to the extracellular matrix^19^. Subsequently, the microenvironment undergoes a transition to an anti-inflammatory stage that involves distinct macrophages in scar resolution^16, 20^. Moreover, macrophages are reported to stimulate cardiomyocyte proliferation during heart development through activation of the epicardium and promote angiogenesis during wound healing^21, 22^. This leaves open the possibility that the different immune responses to cardiac injury may underly the diametrically opposed cardiac and extra-cardiac outcomes between adult mammals and zebrafish.

Here we used unbiased single cell transcriptomics to profile the dynamic immune response to cardiac injury in the heart, blood, liver, kidney, and pancreatic islets of mice and zebrafish. This allowed us to identify analogous immune cell subtypes between the two species along with conserved and disparate responses to cardiac injury within the heart and peripheral organs. Despite similar cardiac macrophage subclusters present in both mice and zebrafish, the response of the two were dramatically different to myocardial injury. Additionally, there were unique macrophage subclusters to each of the species. This study provides support for both differences in macrophage subtypes between species and the largely disparate reactions to heart injury among the analogous macrophage subtypes that may determine healing versus heart failure with systemic inflammation.

## Results

### Single cell transcriptomic dissection of local and peripheral cellular responses to myocardial infarction in mouse

To interrogate the multiorgan response to myocardial infarction (MI), we used 10-12 week old mice and performed a permanent left anterior descending (LAD) coronary artery ligation or a sham surgery (Fig. 1a). LAD ligation resulted in severe left ventricle (LV) systolic dysfunction with an average left ventricular ejection fraction (LVEF) of 27% accompanied by a thinned LV wall ( ). Tissues were collected at 1, 7, and 30 days post injury (dpi) (Fig. 1a). The 1 dpi time point is representative of an acute MI with the initial proinflammatory phase, while day 7 is part of the subacute phase when the inflammatory response shifts toward a reparative phase^23^. Finally, the 30 dpi time point represents chronic MI with heart failure (HF)^24^. We performed scRNA-Seq on isolated single cells from heart (LV), white blood cells (WBCs), liver, and pancreatic islets at the 3 time points. For the kidney, scRNA-Seq was performed at 7 and 30 dpi. After curating the sequencing data for low read counts and high mitochondrial gene ratios per cell, a combined ∼196,000 quality cells were included in the analyses. The cells were unbiasedly clustered and plotted in a UMAP. Cell type labeling was first done using SingleR and then validated by the expression of known marker genes for each cell type (Fig. 1b and Extended Data Fig. 1d)^25^.

**Fig. 1:**
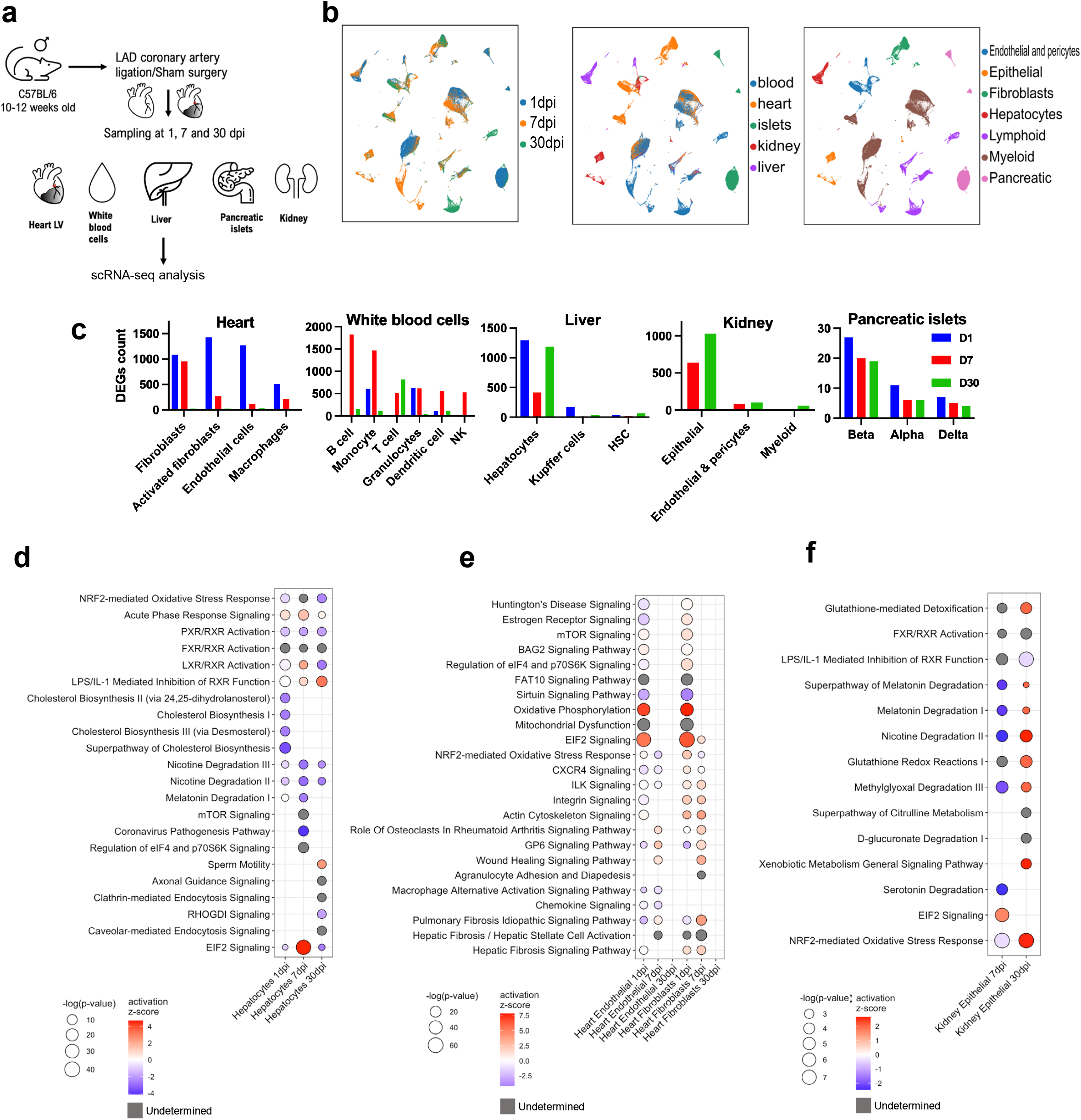
Multiorgan response to MI in mouse by scRNA-Seq. **a,** Experimental outline for mouse LAD ligation and sampling for scRNA-seq analysis. **b**, UMAPs of the 194,315 cells from 41 mice colored by time point (left), tissue of origin (center) and cell type (right). Dpi – days post infarction. **c,** Bar plot of DEGs (MI compared to sham procedure) in mice by tissue and cell type. Bars are colored by time point (days 1, 7 and 30). **d-f,** Dot plots depicting enriched pathways and their statistical significance and activation z-score of hepatocytes (**d**), heart endothelial and fibroblasts (**e**), and kidney cells (**f**) between sham and MI by dpi. Positive z-score predicts activation of the pathway in the MI compared to the sham group. Gray dots denote undetermined activation status.

Across the various tissues, leukocytes yielded the most cells for analysis with over 10,000 cells per sample. Pancreatic islet and heart immune cells were the next most abundant with over 2000 cells, except for heart samples at 30 dpi (Extended Data Fig. 1e). For 30 dpi heart samples, comparatively smaller number of immune cells were sequenced (∼500 cells in the MI vs ∼800 in the sham), indicative of locally resolved inflammation. Smaller numbers of cardiomyocytes were sequenced as we did not employ single nuclei RNA-Seq (snRNA-Seq) and this was not the focus of our study. In the liver, we sequenced over 300 immune cells per sample, except for 30 dpi sham, which contained fewer immune cells (Extended Data Fig. 1e).

Most WBCs were granulocytes at 1 dpi with similar frequencies between sham and MI (Extended Data Fig. 1e). However, at 7 dpi the percentage of monocytes vastly declined in the MI group compared to sham controls. By day 30, T cells were the most abundant WBC in both groups. In the heart, the major immune cell type were myeloid cells at all time points. Granulocytes were remarkably elevated on day 1 in MI (>25% of sequenced cells), and B cells were elevated in MI at 30 dpi, albeit their percentage was modest (<10% of sequenced cells). The scRNA-Seq blood cellular composition results were independently confirmed by flow cytometry in separate cohorts of mice (Extended Data Fig. 1f,g). In the liver, the most abundant immune cells were myeloid in origin, including Kupffer cells. Whereas nearly all (95%) of the immune cells in the liver at day 30 in the sham group were Kupffer cells, they only accounted for ∼20% of the immune cells in the MI group with non-Kupffer myeloid cells and lymphocytes making up the majority of the immune infiltrate. Finally, cellular composition was mostly unchanged with MI in the kidney and pancreatic islets. Most of the kidney cells sequenced were epithelial or endothelial cells while beta cells were the dominant cell type in the pancreatic islets (Extended Data Fig. 1e).

### MI instigates a multiorgan inflammatory response in mouse

To identify which organs and cell types were most strongly perturbed in response to MI, we plotted the number of differentially expressed genes (DEGs) between sham and MI, by organ, cell type, and time point. At 1 dpi, the heart and liver had by far the greatest number of DEGs (Fig. 1c). Over 1000 significant (FDR<0.05 and absolute log_2_(FC)>1) DEGs were detected in heart fibroblasts and endothelial cells and liver hepatocytes. At 7 dpi, the most DEGs were detected in blood B cells, monocytes and T cells. Heart fibroblasts, kidney epithelial cells and hepatocytes all had over 400 DEGs 1 week after MI. By 30 dpi, the most DEGs were detected in hepatocytes and in blood T and B cells and monocytes. Pancreatic islet cells displayed minor transcriptomic alterations following MI (Fig. 1c). To gain insight how each cell type reacted in response to MI, we searched for pathways enriched in the sham and MI groups for each of the organs, cell types, and time points (Fig. 1d-f).

### Persistent inflammation and metabolic changes in hepatocytes after MI

MI resulted in dramatic changes in hepatocyte metabolism. EIF2 Signaling was especially enriched and active on day 7 in the MI (Fig. 1d and Extended Data Fig. 2a). A similar activation pattern was detected for LXR/RXR activation. Acute Phase Response Signaling showed an activation pattern in the MI reflecting the organ’s persistent stress response. By contrast, Melatonin and Nicotine Degradation and PXR/RXR Activation pathways were downregulated in the MI group compared to sham. The NRF2-mediated Oxidative Stress Response pathway was enriched throughout with tendency to be inhibited, especially at 30 dpi. Some pathways were only enriched at one time point. At 1 dpi, cholesterol biosynthesis pathways were inhibited in the MI group compared to controls (Fig. 1d and Extended Data Fig. 2a). At 7 dpi, hepatocytes from MI were enriched in the mTOR and eIF4 and p70S6K signaling pathway with strong inhibition of the Coronavirus pathogenesis pathway. By day 30 after MI, endocytosis pathways such as clathrin and caveolar-mediated endocytosis and RHOGDI Signaling were enriched. Altogether the gene expression and pathway analysis data indicate persistent inflammation and dynamic changes in metabolic pathways in hepatocytes with MI.

### Acute activation and resolution of stress, inflammatory, and fibrotic pathways in heart endothelial cells and fibroblasts after MI

Heart fibroblasts and endothelial cells exhibited a similar enrichment in pathways in response to MI that were strongest at 1 dpi, faded at 7 dpi, and were completely resolved by 30 dpi (Fig. 1e). The EIF2 Signaling and Oxidative Phosphorylation pathways were strongly upregulated at 1 dpi with MI. Mitochondrial dysfunction, Sirtuin signaling (inhibited), Huntington’s disease signaling, Estrogen Receptor Signaling, NRF2 stress, CXCR4, and Integrin signaling pathways were also strongly enriched at 1 dpi in the MI compared to controls (Fig. 1e). These pathways are consistent with the inflammation, organ stress and damage occurring during the acute phase of the MI at the injury site. However, by 7 dpi the transcriptomic landscape had remarkably changed and was representative of myocardial remodeling with Wound Healing and Agranulocyte Adhesion and Diapedesis pathways enriched. Furthermore, pathways related to fibrosis and angiogenesis such as Pulmonary Fibrosis, Hepatic Stellate Cell Activation, and ILK Signaling were activated in the fibroblasts in the MI group compared to sham at 7 dpi with resolution by day 30. Overall, these data suggest that heart endothelial cells and fibroblasts are among the most affected cells in the injury site at day 1 and continue to play a critical role in myocardium remodeling in the first week after injury before returning to baseline at 1 month post infarction.

### Kidney epithelial cell dysfunction after MI

We analyzed kidney cells at 7 and 30 dpi to focus on the longer-term extracardiac effects of MI. At both time points, the top enriched pathway was NRF2-Mediated Oxidative Stress Response, which showed a strong activation pattern at 30 dpi in the MI compared to sham (Fig. 1f). This pathway is aimed at controlling the detrimental effects of excess oxidative stress and reflects potential long-term stress or damage to the kidneys. Many of the other overrepresented pathways were related to the degradation of methylglyoxal, nicotine, serotonin, melatonin, and glutathione and showed an inhibited score at 7 dpi but an activated score at 30 dpi in the MI group compared to controls. The genes common to these pathways *Akr1a1, Cyp4b1,* and *Cyp2e1* were initially downregulated in the MI at 7 dpi and later upregulated at 30 dpi (Extended Data Fig. 2b). This pattern of activation is consistent with early reduced organ function followed by later recovery or compensation after MI. In sum, at both 7 and 30 dpi, kidney epithelial cells in the MI group were enriched in pathways compatible with organ stress and altered function, highlighting the long-term effects of MI on extracardiac organs such as the kidney.

### Chronic systemic inflammation in mice following MI

We found that MI induced a strong transcriptional response in all of the major immune cell types across multiple organs during the acute, subacute, and chronic phases of MI. In blood, B cells had 1827 DEGs at 7 dpi, while T cells had 814 DEGs at 30 dpi (Fig. 1c). Myeloid cells were the most abundant leukocytes in all of the organs we sequenced (heart, liver, kidney and pancreas) at all time points. While the ratio of sequenced parenchymal cells minimally changed, immune cell ratios and absolute counts displayed dynamic changes in response to MI (Extended Data Fig. 1e-g). By scRNA-Seq, Blood B and T cells increased dramatically over time in both sham and MI groups, from close to 0% at 1 and 7 dpi to a majority of WBCs at 30 dpi (Extended Data Fig. 1e,g). This is consistent with previous reports that major surgeries or trauma can cause transient lymphopenia in blood^26^ and considered a side effect of the surgery.

The frequency of blood monocytes decreased from ∼40% at 1 dpi to <20% at 30 dpi in the MI group. However, in the sham group, monocytes spiked to ∼80% of the total WBC at 7 dpi and then decreased to 2% at 30 dpi. By flow cytometry we observed similar frequencies, with lymphocytes increased at day 30 compared to day 7, and myeloid cells following the opposite trend (Extended Data Fig. 1g). In the rest of the analyzed tissues, the most abundant immune cells were myeloid cells. In the heart at 1 and 7 dpi, the MI group had increased numbers of monocytes/macrophages (2800 vs. 2100 at 1 dpi and 3900 vs. 1500 at 30 dpi in MI vs. sham) and granulocytes at 1 dpi (1600 vs 130) compared to sham. By day 30, both sham and MI groups had similar number myeloid cells in the heart. In contrast, liver myeloid cells took up a similar proportion (∼30% sham vs 40% MI) of the liver immune cells at 1 and 7 dpi; however, at 30dpi, the sham was exclusively composed of Kupffer cells whereas the MI group persistently consisted of a diverse set of immune cells such as T, B, and NK cells (Extended Data Fig. 1e). The increased numbers of macrophages and T and B lymphocytes at 30 dpi in the livers suggest unresolved liver inflammation after MI. Similarly, in the kidney we observed >2-fold increase in the percentage of macrophages at 30 dpi in the MI compared to sham (∼12% MI vs. 5% sham). Next, we examined each of the different immune cell subsets to gain further insight into how MI perturbed each over the different phases of injury.

### Lymphocyte immunomodulation with chronic MI

At 1 dpi, very few T and B cells were sequenced, precluding reliable pathway analysis. However, blood T and B lymphocytes were abundant at 7 and 30 dpi (Extended Data Fig. 1e,g). At 7 dpi, T cells displayed enrichment of Oxidative Phosphorylation, Mitochondrial Dysfunction, Actin Cytoskeleton, Integrin Signaling, and Th1 pathway with most of them showing an inhibited score (Extended Data Fig. 3a). At 30 dpi, there were fewer enriched pathways in blood T cells with MI and many related to increased protein translation and metabolic activity (EIF2, Regulation of eIF4 and P70S6K, and mTOR Signaling). The Th1 pathway remained enriched and at 30 dpi showed a tendency towards activation (Extended Data Fig. 3a). Overall, these pathways suggest T cell inhibition early on with MI that shifts towards a Th1 phenotype with chronic MI.

Blood B cells had the most DEGs of any cell type at 7 dpi (Fig. 1c). The top enriched pathways were related to protein translation and metabolic activity such as EIF2 Signaling, Regulation of EIF4 and p70S6K Signaling, mTOR Signaling, and Oxidative Phosphorylation (Extended Data Fig. 3b). At 30 dpi, most of the top pathways enriched in B cells with MI were not seen at 7 dpi, including Primary Immunodeficiency, inhibition of T Cell Receptor Signaling,and CTLA4 Signaling in Cytotoxic T Lymphocytes, suggesting a dampened humoral immune and T cell inhibitory response by B cells in chronic MI compared to control.

### MI triggered a sustained inflammatory response in monocytes

From 1 to 30 dpi, MI induced a gene program in monocytes enriched in adhesion and diapedesis, Macrophage Alternative Activation, IL-10 Signaling, Acute Phase Response, and Pathogen-Induced Cytokine Storm Signaling, suggestive of broad inflammatory activation with heart injury (Fig. 2a). At 7 dpi there was enrichment of EIF2 Signaling, mTOR Signaling, Fc gamma Receptor-mediated Phagocytosis, and Integrin Signaling pathways with many showing an inhibited gene expression pattern (Extended Data Fig. 2c). While at 30 dpi, the Complement System, Antigen Presentation Pathway, and Macrophage Alternative Activation Signaling pathways were upregulated in the monocytes with MI. In summary, our analyses revealed that monocytes exhibited an activated-inhibited-activated pattern at 1, 7 and 30 dpi with MI compared to sham. Intriguingly, some of the inflammatory pathways active on day 1 were active a month later in WBCs even after resolution of inflammation in the heart, indicating persistent monocyte activation with chronic MI. Moreover, the simultaneous detection of pro and anti-inflammatory pathways in monocytes suggests the coexistence of different monocyte subpopulations that may reflect the complex and multifaceted response of these cells to MI.

**Fig. 2:**
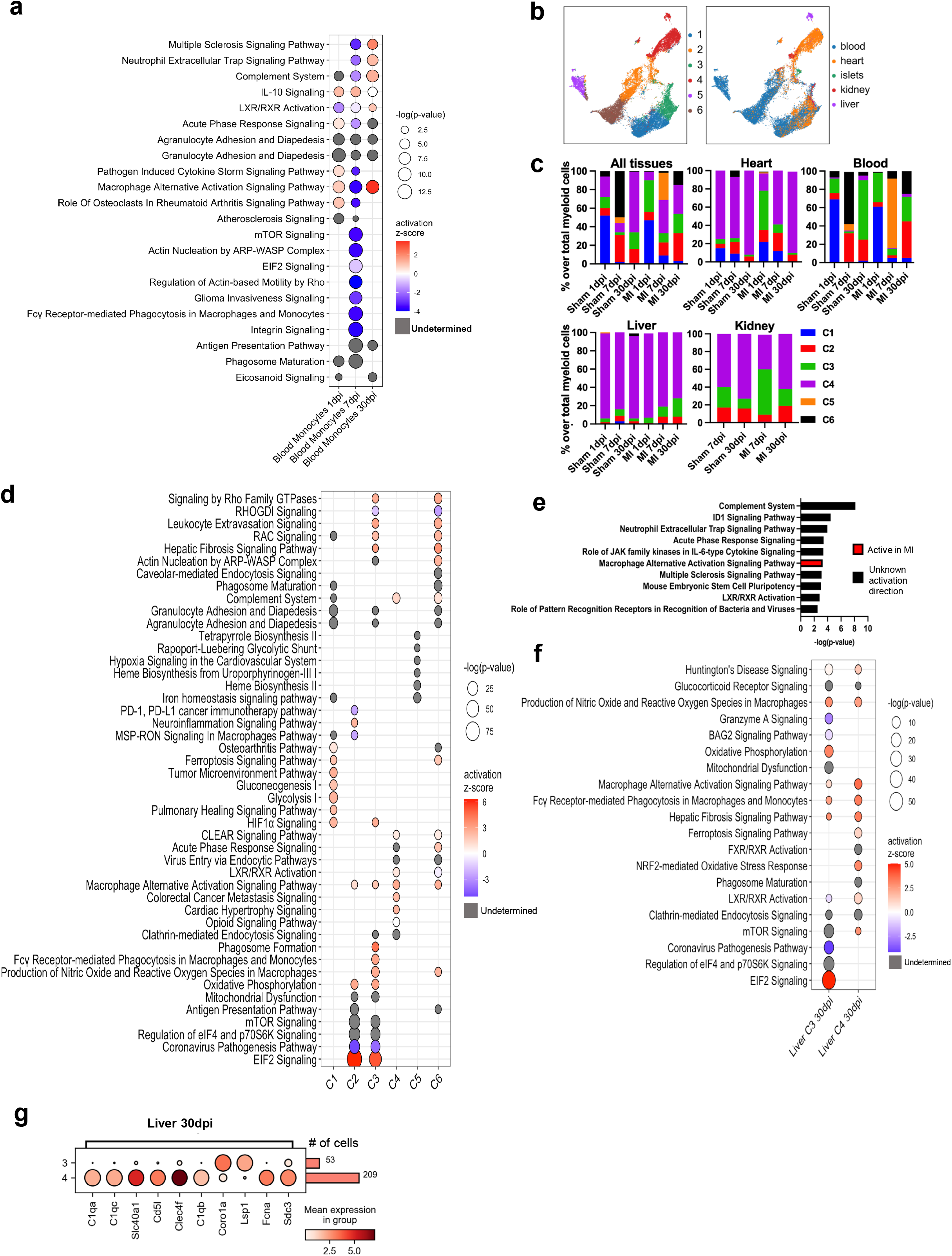
Immune cell response to MI in mouse by single cell transcriptomic analyses. **a,** Dot plot depicting enriched pathways and their statistical significance and activation z-score of monocytes at 1, 7, and 30 days post infarction (dpi). Positive z-score predicts activation of the pathway in the MI compared to the sham group. Gray dots denote undetermined activation status. **b,** UMAPs of all 37,215 sequenced myeloid cells (monocytes and macrophages) reclustered, colored by cluster (left) and tissue of origin (right). **c,** Stacked column plot shows percentage of the 6 murine myeloid subclusters (C1-C6) of total myeloid cells by surgery, time point and tissue. **d,** Dot plot depicting enriched pathways and their statistical significance and activation z-score for each of the myeloid cell subclusters, regardless of tissue of origin, time point or type of surgery. **e,** Bar plot of the pathways enriched in MI compared to sham in cluster 2 blood monocytes at 30 dpi. **f,** Dot plots of the enriched pathways and their statistical significance and activation z-score for the selected myeloid cell subclusters in liver at 30 dpi. In **a**, **d** and **f**, gray dots denote undetermined activation status.

### Distinct monocyte/macrophage subsets in tissues perturbed in response to MI

Monocytes and macrophages constituted most of the sequenced immune cells in tissues and a significant portion of the WBCs in our analysis. Myeloid cells comprise a heterogenous group of cell types, with significant differences and functions depending on their origin and localization^27^. To determine if MI could induce different responses in specific myeloid subtypes, we reclustered all the monocytes and macrophages, excluding granulocytes and dendritic cells, from the different organs, time points, and sham/MI experimental groups to identify macrophage/monocyte subsets. We refer to these as “myeloid clusters”. We identified 6 subclusters of myeloid cells and performed pathway analyses on the top DEGs to ascertain their function (Fig. 2b-d). Cluster 1 myeloid cells were mostly found in the blood and heart at 1 and 7 dpi and enriched in Glycolysis, Gluconeogenesis, Cytokine Storm Signaling, and Adhesion and Diapedesis pathways, suggesting a classical proinflammatory phenotype (Fig. 2d).

Cluster 2 myeloid cells increased in numbers in the blood following 1 dpi in both sham and MI groups and were highly enriched in Antigen Presentation, EIF2, PD-1, and mTOR signaling pathways, suggesting immune modulation and increased RNA translation and metabolic activity in these cells. Cluster 3 cells were present in the blood of both experimental groups but found at elevated numbers in the heart at 1 dpi and kidneys and liver at 30 dpi in the MI group (Fig. 2c). The cluster 3 myeloid cells were enriched in the Fcγ Receptor-Mediated Phagocytosis, Production of Nitric Oxide and Reactive Oxygen Species, EIF2, and Regulation of EIF4 and p70S6K Signaling, and metabolic pathways (Oxidative Phosphorylation, Mitochondrial Dysfunction, and mTOR Signaling) with an activation pattern suggesting high metabolic and phagocytic activity (Fig. 2d).

By contrast, Cluster 4 cells were found in the tissues and not the blood, consistent with a tissue macrophage phenotype (Fig. 2c). Supporting this, cluster 4 macrophages in the liver had high expression of *Clec4f*, a Kupffer cell or liver macrophage marker (Fig. 2g). Cluster 4 macrophages were enriched in LXR/RXR activation, endocytosis, complement, and phagocytic pathways, all of which may suggest tissue homeostatic and clearance functions (Fig. 2d). Myeloid cluster 5 was temporarily present in the blood of the MI group only at 7 dpi. DEGs for Cluster 5 suggest a role for these cells in iron homeostasis and hypoxia with Heme and Tetrapyrrole Biosynthesis and hypoxia pathways enriched. Lastly, Cluster 6 cells represented over half (57%) of the blood monocytes in the sham group at 7 dpi and were overrepresented (25%) in the MI group at 30 dpi when these cells were present at trace amounts (4%) in the sham group (Fig. 2c). Complement System, Leukocyte Extravasation Signaling, Antigen Presentation, and Hepatic Fibrosis were among the top pathways overrepresented in Cluster 6 myeloid cells with an activation pattern suggesting an overall, proinflammatory phenotype. Its top marker gene was *Saa3* (Extended Data Fig. 3c), which is reported to increase monocyte and macrophage cytokine production^28^.

### Cluster-specific myeloid changes in chronic MI

Describing the long-term inflammatory complications of MI in extracardiac organs was the focus of this study. Thus, we centered our attention to persistent differences between sham and MI in the myeloid subpopulations across organs. Compared to the sham, there was a larger representation of cluster 2 in blood (23% sham vs 40% MI) and cluster 3 in the liver (3% sham vs 20% MI) at 30 dpi (Fig. 2d). We hypothesized that those cells could be driving the MI-derived persistent inflammation. To interrogate their potential function, we performed IPA on these cell subpopulations in blood and liver at 30 dpi, comparing each cluster against the rest of the myeloid cells of the tissue, regardless of sham or MI surgery (Fig. 2e,f). Cluster 2 blood monocytes in the MI group were enriched in complement system, ID1 signaling, neutrophil activation, and macrophage alternative activation pathways compared to sham controls, suggesting immune activation of this subset with chronic MI (Fig. 2e).

In the liver, cluster 3 myeloid cells exhibited signs of increased protein translation (EIF2) and metabolic activity (Oxidative Phosphorylation) in the MI compared to sham group at 30 dpi (Fig. 2f). However, Granzyme A and Coronavirus pathogenesis pathways were inhibited in the MI group, suggesting altered killing activity. Cluster 4 was the most numerous myeloid subpopulation in the liver and mainly constituted of Kupffer cells expressing the marker gene *Clec4f* (Fig. 2g). Kupffer cells are the first responders of liver injury and play important roles in liver disease, including the recruitment of bone marrow derived monocytes, regulation of hepatocyte proliferation and fibrosis. In addition, Kupffer cells play several functions in tandem with hepatocytes, such as iron recycling and cholesterol homeostasis^29^. Our data indicate that MI resulted in liver damage and hepatocyte dysfunction at 30 dpi. Most of the enriched pathways with MI in Cluster 4 cells were associated with macrophage activation and phagocytosis and showed an activation pattern compared to sham (Fig. 2f). Hepatic Fibrosis signaling was also active in the MI, suggesting chronic liver injury. To compare the 2 most abundant macrophages in the liver at day 30, we also plotted the DEGs between liver clusters 3 and 4. The top DEGs (by FDR) were three genes of complement C1 (*C1qa, C1qc,* and *C1qb*) and other genes related to complement system activation (*Cd5l*^30^ and *Fcna*^31^), which were all higher in cluster 4 than 3 (Fig. 2g). The gene coding for ferroportin (*Slc40a1*), a protein involved in releasing iron to the extracellular space, was also elevated in cluster 4. Moreover, immunohistochemical (IHC) staining for macrophages revealed double the number of hepatic macrophages in the MI group compared to sham at 30 dpi, validating the scRNA-Seq data (Extended Data Fig. 3d). Altogether, our data suggest altered liver macrophage composition and gene expression changes signaling increased inflammation and tissue damage in the MI group compared to controls.

### Cardiac and systemic responses after heart injury in zebrafish

For comparisons between regenerative and non-regenerative species, we performed a similar scRNA-seq of zebrafish cells after heart cryoinjury. We isolated cells from the heart, liver, pancreas, and whole kidney marrow (WKM) at 1 and 7 days post heart cryoinjury (dpci) together with the uninjured control (Fig. 3a). These 2 time points represent acute inflammation (1 dpci) and regenerative stages (7 dpci), respectively^15^. A total of 70,783 cells from 2 replicates of 4 organs and 3 treatment groups (uninjured or Ctrl, 1 dpci, and 7 dpci) passed quality control filters and were included in the analyses (Fig. 3b and Extended Data Fig. 4). Unbiased clustering yield 16 clusters with the major cell types annotated by known marker genes (Fig. 3c-e and Extended Data Fig. 5a). As expected, heart cells make up the most significant changes in cell type fractions (Extended Data Fig. 5b, c). Our analyses included 3613, 5613, and 4475 cells for the uninjured (Ctrl), 1 dpci, and 7 dpci heart samples, respectively when 2 replicates were combined. Erythrocytes, expressing *cahz* and *alas2*, constitute about 60% of the total cells (Extended Data Fig. 5a-c). The uninjured sample contains about 11% CMs (*myl7*^+^*tnnt2a*^+^), 7% macrophages (or monocytes, *c1qa*^+^), 5% NK or T cells (*zbtb32*^+^*il2rb*^+^ or *lck*^+^*zap70*^+^), 3% thrombocytes (*itga2b*^+^), 2% B cells (*cd37*^+^), and 1% neutrophils (*mpx*^+^). The remaining cell types, including epicardial cells (*tcf21*^+^), endocardial or endothelial cells (*kdrl*^+^), and mural cells (*pdgfrb*^+^), contribute about 14% of all cells, which are labeled as the Mixed cluster 5 (Fig. 3c-e and Extended Data Fig. 5). The macrophage and neutrophil populations sharply expanded to 26% and 6% of all cells at 1 dpci, respectively, indicating an acute inflammatory response. By contrast, the combined number of CMs and Mixed cells was down to about 3%. By 7 dpci, the number of NK/T cells and B cells increased to 8% and 3%, respectively, when macrophages were down to 13% and neutrophils to 1% (Extended Data Fig. 5). These observations match previous reports that that macrophages and neutrophils domine the acute inflammation stage while NK/T cells take on activity during the regeneration stage^15, 16^

**Fig. 3:**
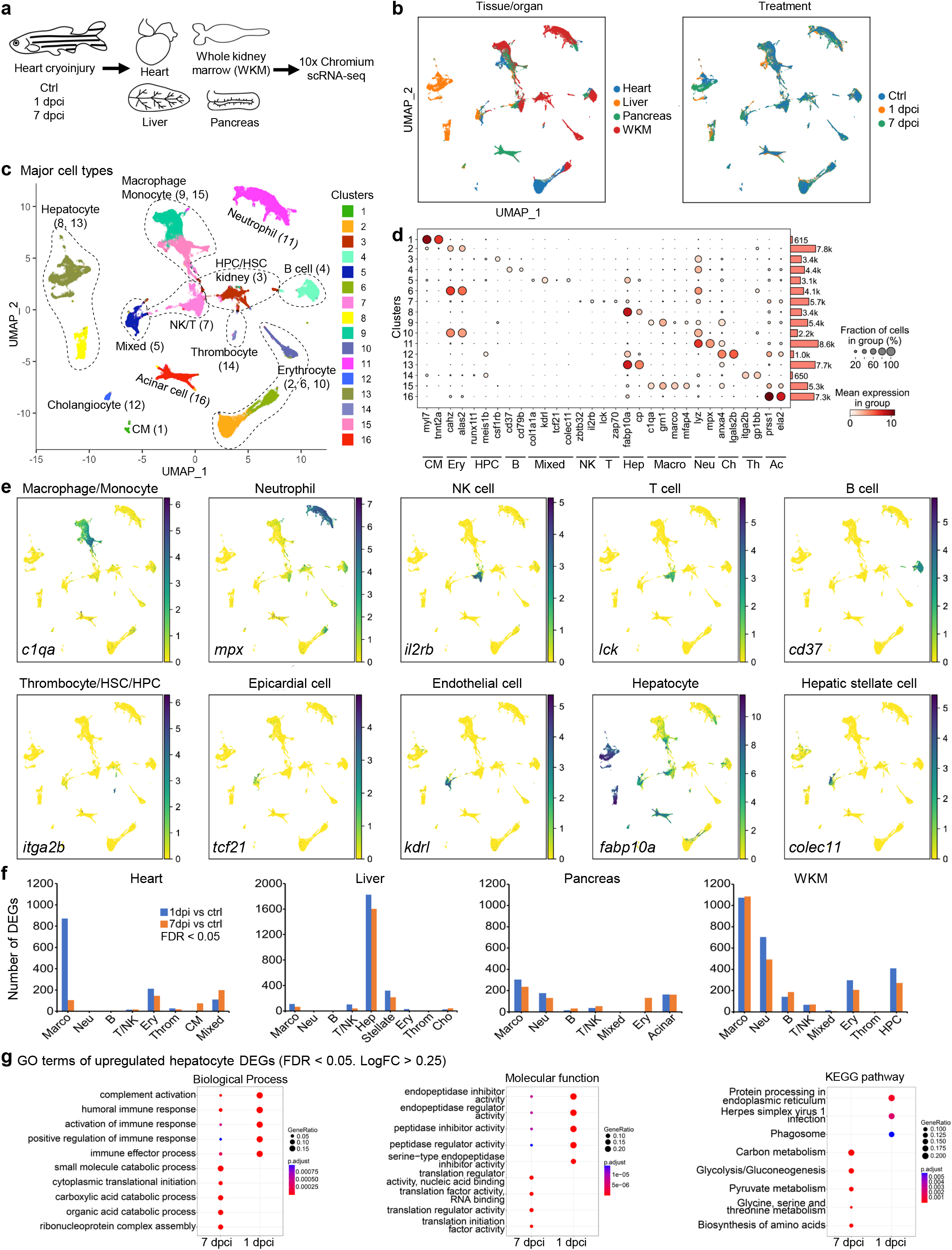
Parenchymal cell response to cryoinjury in zebrafish by scRNA-Seq. **a,** Experimental outline for cryoinjury and tissue collection. **b**, UMAPs of QC-filtered 70,783 cells showing tissue of origin (left) and treatment time points (right). WKM, whole kidney marrow; Ctrl, uninjured. **c**, A UMAP showing 16 clusters with major cell types outlined and labels. Numbers in brackets show clusters. **d**, Dot plot depicting the abundance and expression magnitude of individual marker genes across cells of different clusters. The dot size represents the fraction of cells with at least one UMI of the specified gene. Bars and numbers on the right show cell numbers in each cluster. Major cell types are labeled at the bottom. CM, cardiomyocyte; Ery, erythrocyte; HPC, hematopoietic stem or progenitor cell; Hep, hepatocyte; Macro, macrophage; Neu, neutrophil; Ch, cholangiocyte; Th, thrombocyte; Ac, acinar cell. **e**, UMAPs showing expression patterns of selected marker genes. **f**, Bar plots showing numbers of differentially expressed genes (DEGs) in major cell types from each organ. FDR < 0.05. **g**, Biological process (left), molecular function (middle), and KEGG pathways (right) enriched for the upregulated DEGs (FDR < 0.05, LogFC > 0.25) in hepatocytes at 1 and 7 dpci. The gene ratio is indicated by the dot size and the significance by the color of the dot.

For the liver, cell-type fractions are largely unchanging across samples. We analyzed 7163 (Ctrl), 3354 (1 dpci), and 5097 (7 dpci) cells, of which 70% on average are hepatocytes (*fabp10*^+^*cp*^+^, Fig. 3c-e and Extended Data Fig. 5b, c). Cholangiocytes (*anxa4*^+^*lgals2b*^+^) and stellate cells (*colec11*^+^, in the Mixed cluster 5) make up about 16%. The remaining are 6% macrophages, 6% NK/T cells, and 2% erythrocytes. For the pancreas, 6043 (Ctrl), 4966 (1 dpci), and 5137 (7 dpci) cells are included in the analyses. The zebrafish pancreatic islets and exocrine lobules tightly adhere to intestinal and hepatic tissues. Our dissection method may not be ideal for isolating the islet cells (< 2% of the total, included in the Mixed cluster 5), and the samples likely contain some non-pancreatic cells. The primary cell type fractions in the Ctrl sample are 44% acinar cells (*prss1*^+^*ela2*^+^), 27% macrophages, 9% neutrophils, 9% NK/T cells, and 5% B cells (Fig. 3c-e and Extended Data Fig. 5b, c). The acinar cell fraction increases to 55% at 1 dpci but goes down 38% at 7 dpci. Another major change is the macrophage ratio, which decreases to 19% at 1 dpci and 17% at 7 dpci. The NK/T cell and neutrophil fractions are relatively unchanged at 1dpci but increase to 14% and 13% by 7 dpci, respectively. The WKM samples yielded the most cells that passed the QC, with 7379 (Ctrl), 8344 (1 dpci), and 9608 (7 dpci) cells analyzed. The main cell fractions in the uninjured sample (Ctrl) are 21% neutrophils, 20% macrophages, 20% erythrocytes, 13% hematopoietic stem or progenitor cells (HPC, *runx1t1*^+^*meis1b*^+^*csf1rb*^+^), 14% B cells, and 11% NK/T cells (Fig. 3c-e and Extended Data Fig. 5). The neutrophil population expanded to 32% at 1 dpci and 23% at 7dpci. By contrast, the macrophage fraction shrunk to 15% at 1 dpci and 14% at 7 dpci. The erythrocyte fraction shrunk to 18% at 1 dpci before increasing to 27% by 7 dpci. The numbers of the remaining cell types are largely constant across samples.

We next analyzed the cell numbers of major immune cells (such as macrophages, neutrophils, NK/T cells, and B cells) across all samples. These 29,432 immune cells consist of 36% macrophages, 29% neutrophils, 20% NK/T cells, and 15% B cells (Extended Data Fig. 6a). At 1 dpci, while the number of macrophages sharply increased in the heart, making it the most abundant source, macrophages in other organs decreased synchronously (Extended Data Fig. 6b). It is possible that macrophages from distal organs translocate to the heart upon heart injury to engage acute immune responses. By contrast, the numbers of neutrophils in the heart and WKM increased concurrently at 1 dpci. Interestingly, minor changes in the abundance of immune cells in the liver were detected (Extended Data Fig. 6b), suggesting that the liver likely has minimal immune responses upon heart injury. In summary, compared to the heart samples, cell fractions in the non-cardiac organs are relatively stable upon heart injury, with some fluctuation in immune cell numbers. This may reflect a systemic motivation of immune cells that may relocate to the damaged heart tissue.

### Metabolic but not fibrotic responses in the zebrafish hepatocyte

To better evaluate the systemic responses, we quantified the number of differentially expressed genes (DEGs, FDR < 0.05) at 1 and 7 dpci compared to the uninjured control. Transcriptional changes of cardiac cells upon heart injuries have been intensively studied. Here we focus on systemic changes. Among the main cell types, the most drastic changes were detected from liver hepatocytes, with > 1,600 DEGs at both 1 and 7 dpci (Fig. 3f). To gain insight into how hepatocytes respond to cardiac injury, we performed Gene Ontology (GO) enrichment analysis using the upregulated DEGs. As shown in Fig. 3g, the most enriched GO terms at 1 dpci are complement activation, immune response activation, and endopeptidase related activity, suggesting systemic inflammatory responses at 1 dpci. By 7 dpci, the top terms are metabolic pathways such as carboxylic acid metabolic process, pyruvate metabolism, and glycolysis. Thus, unlike the fibrotic responses in the mouse liver upon MI, zebrafish hepatocytes likely employ enhanced metabolic activities to support heart regeneration at 7 dpci.

### Systemic adaption and transient cardiac inflammatory response in zebrafish macrophages

Besides hepatocytes, we observed transcriptional response in the major immune cell types across organs, with macrophages and neutrophils having the most changes (Fig. 3f). In heart macrophages, we detected ∼800 DEGs (FDR < 0.05) at 1 dpci and 107 DEGs at 7dpci (Fig. 3f), indicating a decreased activity at 7 dpci. Similarly, liver macrophages have 109 DEGs at 1 dpci and 65 at 7 dpci, while pancreatic macrophages have 303 DEGs at 1 dpci and 235 at 7 dpci. By contrast, WKM macrophages show continued drastic activity with ∼ 1000 DEGs at both 1 and 7 dpci (Fig. 3f). This is also true for WKM neutrophils with 703 and 493 DEGs at 1 and 7 dpci, respectively. Pancreatic neutrophils are the second most active after WKM neutrophils, with 176 DEGs detected at 1 dpci and 130 at 7 dpci. In comparison, heart neutrophils are less active, with only 4 and 0 DEGs detected at 1 and 7 dpci, respectively, likely caused by the low number of isolated neutrophils in our heart samples (Extended Data Fig. 6b). Few neutrophils were isolated from liver samples as well. Among other major immune cells, B cells are mostly isolated from the WKM samples, with 142 DEGs detected at 1 dpci and 186 at 7 dpci. This is followed by pancreatic B cells, with 14 DEGs detected at 1 dpci and 32 at 7 dpci. For NK/T cells, liver is the most active organ, with 101 DEGs at 1 dpci and 39 at 7 dpci, followed by WKM NK/T cells, with 66 DEGs at 1 dpci and 69 at 7 dpci. Pancreatic NK/T cells have 37 DEGs at 1 dpci and 53 at 7 dpci, while heart NK/T cells have only 15 DEGs at 1 dpci and 16 at 7 dpci. Thus, we observed systemic immune response in both the injured heart and distal organs and demonstrated drastic activities in macrophages. While cardiac inflammation resolves by 7 dpci, systemic adaption seems to continue from at 7 dpci.

### Subclusters of zebrafish macrophages

Our initial clustering result indicated 2 macrophage or monocyte subpopulations in zebrafish (clusters 9 and 15, Fig. 3c). To further interrogate the heterogeneity and dynamics of zebrafish macrophages after heart injury, we performed sub-clustering of macrophages from all 4 organs. Seven subpopulations were identified with some clusters showed tissue specificity (Fig. 4a). Clusters 1 and 2 are predominantly derived from the WKM, constituting 21% and 14% of all macrophages, respectively (Fig. 4b, c). Cluster 3 (8% of the total) is primarily pancreatic, while cluster 4 (9% of the total) is primarily from the heart (Fig. 4b, c). Cluster 5 is the least subpopulation with only a 2% share. Cluster 6 (31%) is the most abundant, constituting 31% and presenting in all 4 organs. Similarly, cluster 7 has no organ specificity and contributes 15% of all macrophages. Interestingly, four subclusters (1, 3, 6, and 7) were enriched in the uninjured control, with their numbers synchronously decreasing at 1 dpci before bouncing back at 7 dpci (Fig. 4d, e). By contrast, the number of clusters 4 and 5 drastically increased at 1 dpci, but quickly dropped to the uninjured level at 7 dpci, indicating the transiently expanded macrophage subsets in the heart. Cell number of cluster 2 also increased at 1 dpci but maintained at a relatively high level by 7 dpci (Fig. 4e).

**Fig. 4:**
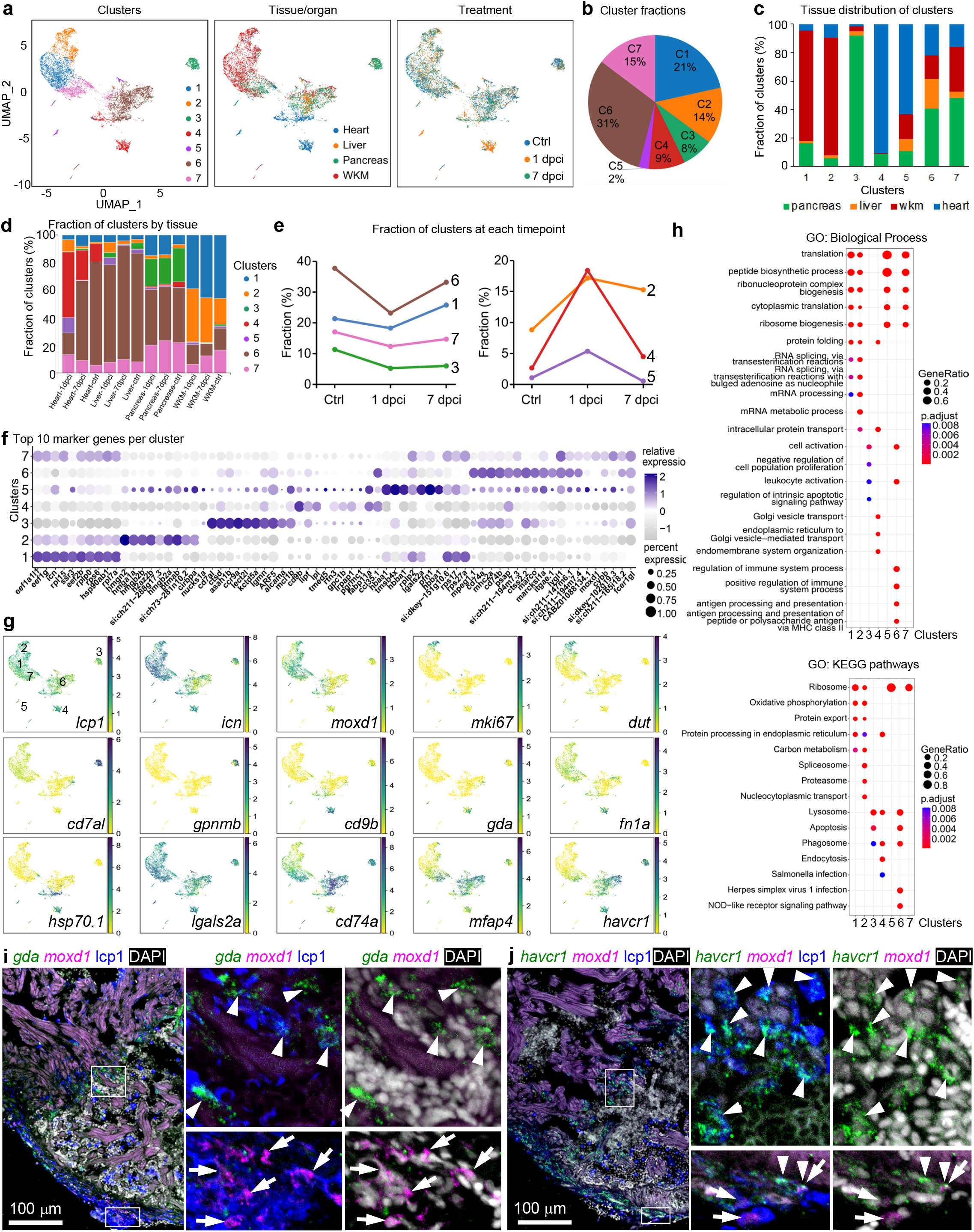
Macrophage/monocyte response to cryoinjury in zebrafish. **a,** UMAPs of 10,709 QC-filtered macrophage/monocytes re-clustered, colored by subcluster (left), tissue of origin (middle), and treatment group (Ctrl, 1 dpci, or 7 dpci; right). **b,** Fractions of each sub-cluster over total macrophages and monocytes. **c,d,** Tissue distribution of each macrophage/monocyte subcluster grouped by cluster (**c**) or tissue and treatment group (**d**). **e,** Dynamics of subcluster fractions over total macrophage/monocyte in each treatment group (Ctrl, 1 dpci, or 7 dpci) across three groups. **f,** Dot plot showing the top 10 marker genes of each subcluster. The dot size represents the fraction of cells with at least one UMI of the specified gene. **g,** UMAPs showing expression patterns of selected marker genes. **h,** Enriched biological process terms (top) and KEGG pathway (bottom) for the subcluster markers (mean AUC > 0.6). The gene ratio is indicated by the dot size and the significance by the color of the dot (P < 0.05). **i**, Images of heart cryosection showing HCR staining signals of *gda* and *moxd1* at 1 dpci in green and magenta, respectively. Anti-lcp1 staining is shown in blue. DAPI staining is shown in white. The framed regions are enlarged to show details on the right with different channel combinations. Arrowheads and arrows indicate representative *gda*^+^lcp1^+^ and *moxd1*^+^lcp1^+^ cells, respectively. Scale bar, 100 μm. **j**, Images of heart cryosection showing HCR staining signals of *havcr1* and *moxd1* at 1 dpci in green and magenta, respectively. Anti-lcp1 staining is shown in blue. DAPI staining is shown in white. The framed regions are enlarged to show details on the right with different channel combinations. Arrowheads and arrows indicate representative *havcr1*^+^lcp1^+^ and *moxd1*^+^lcp1^+^ cells, respectively. Scale bar, 100 μm.

We next check cluster markers to interrogate their functions. The WKM-dominating cluster 1 has enriched expressions of *icn*, *eef1g*, *s100a10b*, *rpl15*, *mrc1b*, and *moxd1*, while cluster 2 is enriched with histone modification and chromatin regulatory genes such as of *hmgn2*, *h2az2b*, *mki67* and *dut* (Fig. 4f, g). Cluster 2 also shares most of the cluster 1 marker genes at a relatively lower expression level. GO term analysis results indicate enriched processes in translation, peptide biosynthetic, and ribonucleoprotein complex biogenesis in both clusters 1 and 2 (Fig. 4h). Therefore, these are the macrophages or monocytes in WKM, and cluster 2 cells are likely the proliferating macrophages or monocytes that replenish the macrophage pool in supporting heart regeneration. The pancreatic cluster 3 showed relatively high expression of *cd7al*, *asah1b*, *ctsd*, *ccr9a*, *lect2l*, *lgmn*, and *gpnmb* (Fig. 4f, g). Notable enriched GO terms in this cluster are negative regulation of cell proliferation, leukocyte activation, and regulation of intrinsic apoptotic signaling pathway (Fig. 4h). The heart-dominated cluster 4 has enriched expression of *cd9b*, *lipf*, *gpc1b*, *fabp11a*, *npc2*, *gda*, *fn1a*, *g0s2*, and *hsp70.1* (Fig. 4f, g). Enriched processes are protein or vesicle transport, protein folding, and endomembrane system organization. Enriched KEGG pathways in cluster 4 are phagosome, lysosome, and endocytosis, suggesting active phagocytosis (Fig. 4h). Cluster 5 has enriched expressions of *hbaa1*, *tmsb4x*, *lgals2a*, *anxa2a*, and *fth1a*, with the top GO terms in translation, peptide biosynthetic process, cytoplasmic translation, and ribosome biogenesis (Fig. 4f-h). Cluster 6 has high expressions of standard macrophage markers such as *mpeg1.1*, *cd74a*, *mhc2a*, *marco*, *mfap4*, and *havcr1*. The top enriched GO terms are mostly regulations of immune system process, antigen processing and presentation, leukocyte activations, and KEGG pathways lysosome, phagosome, and apoptosis (Fig. 4f-h). Top cluster 7 markers include *lygl1*, *si:ch211-147m6.1*, *moxd1*, and *cybb* with enriched GO terms in translation, peptide biosynthetic process, and ribosome biogenesis. Although clusters 6 and 7 are present in all 4 organs, cluster 7 is mostly homeostatic and similar to the WKM clusters 1 and 2, while cluster 6 is pro-inflammatory and participates in phagocytosis. Thus, we predicate that cluster 6 is the resident macrophage population within each organ, and cluster 7 consists of circulating macrophages and/or monocytes.

To better understand functions of these subclusters in the injury heart, we performed hybridization chain reaction (HCR) staining of a few highly specific markers on heart cryosections collected at 1 dpci, together with antibody staining against the pan-leukocyte marker L-plastin (i.e, lymphocyte cytosolic protein 1, lcp1)^32, 33^. As shown in Fig. 4i, expression of the cluster 4 marker *gda* is enriched in macrophages in the boundary zone of the wound. By contrast, the cluster 1, 2, and 7 enriched *moxd1* is mostly on the ventricular surface of the wound (Fig. 4i). In addition, the cluster 3, 4, and 6 enriched *havcr1* is expressed in macrophages both in the boundary zone and ventricular wall at 1 dpci, with some overlap with *moxd1* expression on the ventricular surface (Fig. 4j). These expression patterns suggest that cluster 4 contains boundary zone macrophages while clusters 1, 2, 3, 6, and 7 are mostly on the ventricular surface.

### Shared and distinct cellular and molecular pathways in the dynamic immune response between regenerative and profibrotic myocardial injury models

Given that the immune system plays a critical role in determining the outcome of the cardiac and systemic reaction to myocardial injury, we sought to compare and contrast the immune response between mouse and zebrafish. At 1 and 7 dpi, the monocytes and macrophages showed the largest number of DEGs in zebrafish in comparison to neutrophils and T and B lymphocytes (Fig. 3f). In mice, monocytes and macrophages were amongst the cells with the greatest number of DEGs across the 1 and 7 dpi time points (Fig. 1c). We thus focused our analyses on the myeloid cells (monocytes and macrophages) as they were the most numerous immune cell type and showed dynamic changes with respect to injury across the two species. Macrophages also play critical roles in heart regeneration^18, 19, 34^ First, we remapped all the monocytes and macrophages from both mouse and zebrafish and plotted them on a UMAP (Fig. 5a-c). To compare the 6 myeloid clusters in mice (mm) to the 7 myeloid clusters in zebrafish (dr), we applied the Self-Assembling Manifold mapping (SAMap) algorithm to map single-cell transcriptomes between different species^35^. We found 3 macrophage cluster pairs with high inter-species alignment scores: mm1-dr4, mm4-dr6, and mm6-dr3 (Fig. 5d). mm1-dr4 are injury responsive monocytes/macrophages that are robustly increased in the heart following myocardial injury. The mm4-dr6 pair represents tissue resident macrophages that are present in the heart and other organs examined while the mm6-dr3 myeloid pair was mostly found in the blood in mice and the whole pancreas in zebrafish. Two zebrafish clusters (dr5 and dr7) did not show significant similarity to any mouse myeloid cluster while mouse cluster mm5 did not align with zebrafish. To understand the similarities between these analogous myeloid clusters, we assessed the marker genes for each cluster and performed pathway analyses. Leukocyte migration, pyruvate metabolism, vesicle organization, hypoxia, and autophagy were among the top shared overrepresented pathways for mm1 and dr4 myeloid subclusters (Fig. 5e). The mm4-dr6 pair had the highest number of top 10 shared overrepresented pathways, which related to the complement pathway, humoral immunity, cell junction disassembly, and synapse formation (Fig. 5f). mm6-dr3 had fewer shared pathways that included neutrophil degranulation, immune regulation, and antigen presentation (Fig. 5g). Collectively, these analyses indicate that there are analogous myeloid clusters between mouse and zebrafish that may share similar functions. In addition to the transcriptional differences in the analogous myeloid clusters, there are different myeloid clusters that are distinct to each species.

To examine shared and divergent species transcriptional response to myocardial injury in the myeloid cluster pairs, we focused on the heart where macrophages and infiltrating monocytes are abundant and important in tissue repair as well as the blood, which is an indicator of the systemic inflammatory response and an important source of infiltrating monocytes. We assessed transcriptional changes induced by myocardial injury (fold change compared to control) in myeloid pairs in tissues and time points where they are jointly found: mm1-dr4 and mm4-dr6 at days 1 and 7 in the heart and mm6-dr3 in the blood (Fig. 6 and Extended Data Fig. 7). Focusing just on ortholog genes from the two species, the number of genes that were concordantly upregulated or downregulated in the myeloid cluster pairs was very small compared to those genes that were discordantly transcribed in response to myocardial injury (Fig. 6a, c, e, g and Extended Data Fig. 7a, b). In the heart, both mm1-dr4 and mm4-dr6 had larger number of DEGs at day 1 compared to 7. Pathway analysis of DEGs in the heart at day 1 for mm1-dr4 revealed that genes upregulated in zebrafish but not in mouse were related to neutrophil degranulation and catabolism of nucleoside triphosphate and amino sugars (Fig. 6b). Gene pathways that were overrepresented in mm1 but not in dr4 with injury at day 1 were in general positive for translation, gene expression, and cellular protein targeting while day 7 pathways were neutrophil degranulation, apoptosis regulation, and response to thyroid hormone (Fig. 6d). Comparing the mm4-dr6 clusters in the heart, there was a large transcriptional response induced with injury in the mouse but not zebrafish at day 1 (429 genes) that went down dramatically by day 7 (55 genes) (Fig. 6e, g). At 1 day of injury, the overrepresented pathways in mm4 but not dr6 were categorized by translation, gene expression, macromolecule biosynthesis, and protein localization, all of which suggest robust cellular activation (Fig. 6f). At day 7, mm4 but not dr6 injury response genes were enriched in neutrophil degranulation, cellular response to thyroid hormone, and proteolysis pathways (Fig. 6h). In contrast, in the blood the difference in heart injury response between mm6 and dr3 was negligible at day 1 and most pronounced at day 7 with 238 genes that were upregulated in mm6 but decreased or not changed in dr3 and another 272 genes that were decreased in mouse but not changed in zebrafish (Extended Data Fig. 7a, b). The genes induced by injury at day 7 in mm6 but not in dr3 most strongly fell in pathways related to protein targeting, translation and peptide biosynthesis, and mRNA catabolism (Extended Data Fig. 7c). Whereas the genes that were decreased in mm6 but not in dr3 were enriched in pathways such as neutrophil degranulation, interferon-γ signaling, actin cytoskeleton organization, and foam cell differentiation (Extended Data Fig. 7d). Taken together, these data suggest that the analogous myeloid pairs in mice and zebrafish react differently to heart damage that may underlie the disparate outcomes of fibrosis and regeneration, respectively.

**Fig. 5:**
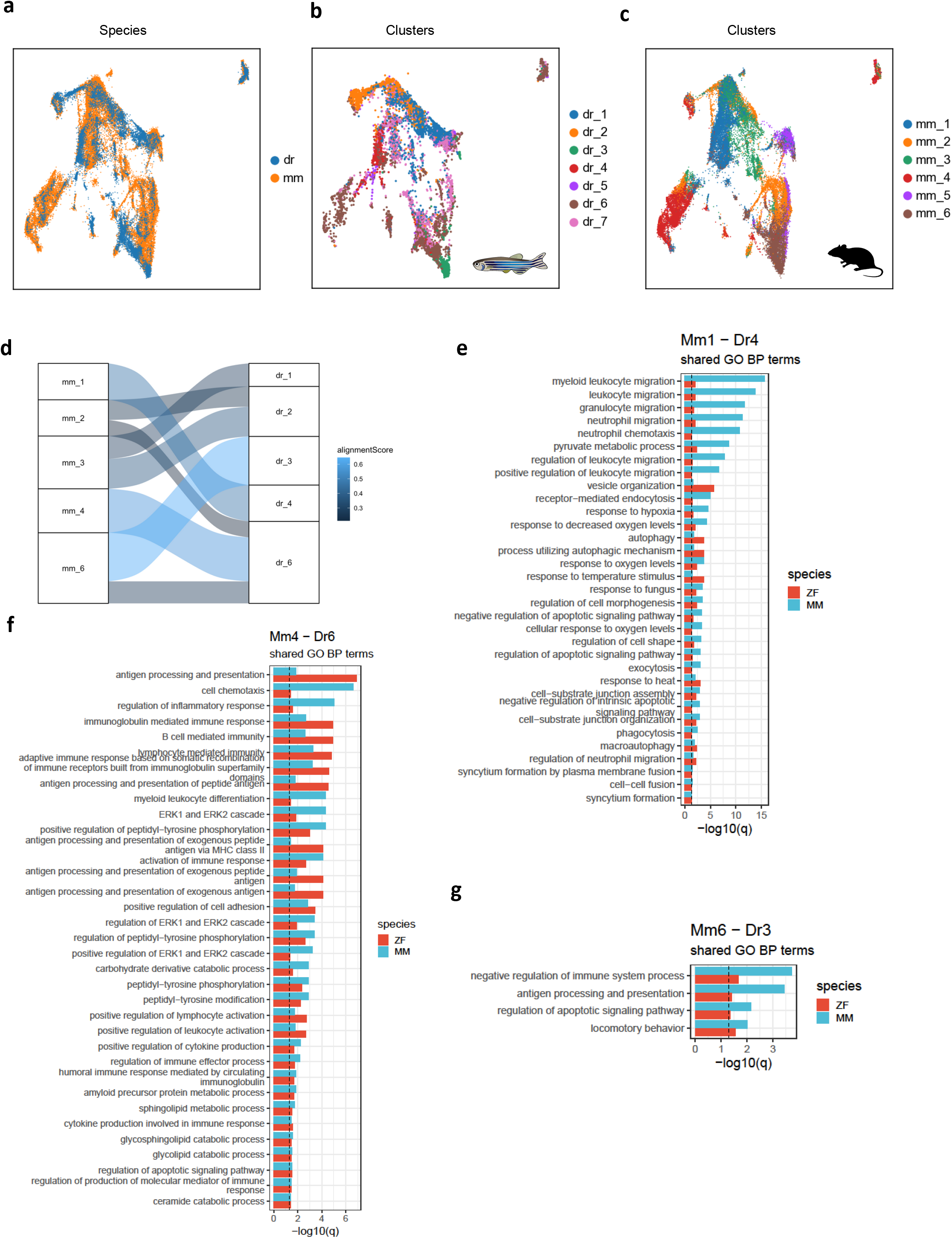
Interspecies similarities and differences in myeloid subsets. **a-c,** UMAPs of myeloid cells from the mouse (37,215 cells) and zebrafish (10,709 cells) plotted as total (**a**), zebrafish (**b**) and murine (**c**) clusters. **d,** SAMap alignment of the mouse (mm) and zebrafish (dr) myeloid subclusters. Edges with alignment scores less than 0.2 were omitted. **e-g,** Shared enriched biological process terms (GO BP) between mm1-dr4 (**e**), mm4-dr6 (**f**), and mm6-dr3 (**g**) clusters based on subcluster markers for each species (mean AUC > 0.6). Only terms with an adjusted p < 0.05 in both species were included.

**Fig. 6:**
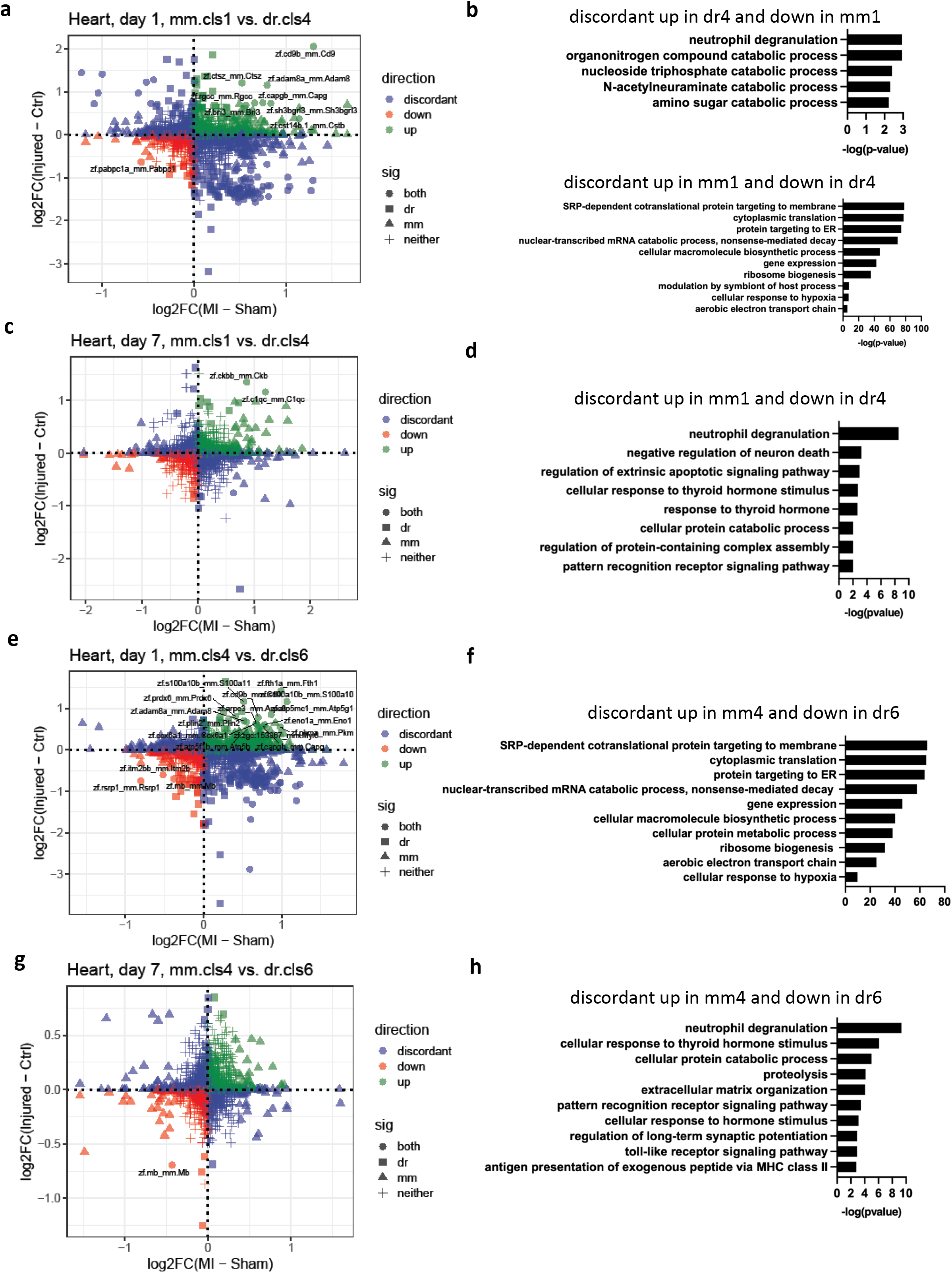
Differential transcriptional response to cardiac injury in myeloid subsets between species. **a**, Scatterplot of the fold changes in murine and zebrafish genes with cardiac injury compared to control in mm1 and dr4 cells at 1 dpi (FDR<0.10 and |log2FC|>0.25). **b,** Enriched biological process terms of the DEGs that are discordant between mm1 and dr4 cells in **a**. **c,** Scatterplot of the fold changes in murine and zebrafish genes with cardiac injury compared to control in mm1 and dr4 cells at 7 dpi (FDR<0.10 and |log2FC|>0.25). **d,** Enriched biological process terms of the DEGs that are discordant between mm1 and dr4 cells in **c**. **e,** Scatterplot of the fold changes in murine and zebrafish genes with cardiac injury compared to control in mm4 and dr6 cells at 1 dpi (FDR<0.10 and |log2FC|>0.25). **f,** Enriched biological process terms of the DEGs that are discordant between mm1 and dr4 cells in **e**. **g,** Scatterplot of the fold changes in murine and zebrafish genes with cardiac injury compared to control in mm4 and dr6 cells at 7 dpi (FDR<0.10 and |log2FC|>0.25). **h,** Enriched biological process terms of the DEGs that are discordant between mm1 and dr4 cells in **g**. In **b, d, f,** and **g,** pathways with a -log_10_(p-value) > 1.3 were considered significantly enriched.

The inter-species differences in the monocyte and macrophage response to heart injury observed by scRNA-Seq was striking. We sought to use an orthogonal system for validation that does not require lysing the cells for analysis and that can be assessed by other labs in their models. In the mouse myeloid subclusters, we identified and plotted marker genes that are commonly used for flow cytometry analysis to characterize macrophages. We plotted macrophage lineage and subtype genes for cardiac macrophages at 1 dpi for the mouse clusters (1-4) present in the heart (Extended Data Fig. 8a). *H2-Aa* was highest in mm2 cells while *Ly6c2* and *Mrc1* were most abundantly expressed in mm3 and mm4 cells, respectively. Flow cytometry of digested cardiac cells allowed us to distinguish macrophage subsets based on cell surface protein expression of these macrophage genes (Extended Data Fig. 8b). Mice with sham and MI at 1 dpi had different fractions of cardiac macrophage subsets, corroborating the scRNA-Seq analyses (Fig. 7a and Extended Data Fig. 8c, d). CD206^+^ macrophages are predicted to be mm4 cells and are 3-fold lower in the MI compared to sham. On the other hand, CD206^-^Ly6c^low^ macrophages would be the equivalent to mm1 macrophages and were about 50% more frequent in the MI than sham group. This further permitted us to use fluorescent activated cell sorting (FACS) to enrich for mm1 cells and assess for transcriptional responses to MI within the different cardiac macrophage subsets. In CD206^-^Ly6c^low^ macrophages, *Fth1* was elevated in the MI group compared to sham (Fig. 7b). However, *Ahnak* and *Psap*, two of the genes that were induced at 1 dpci in the dr4 macrophage subset were not changed with MI in CD206^-^Ly6c^low^ cardiac macrophages (Fig. 7b, c). We performed HCR staining of *cd9b* and *fn1a* on zebrafish heart sections. As shown in Fig. 7e, these two genes are induced in macrophages around the injury site at 1 dpci but were not detectable in the absence of heart injury. Similarly, as we have shown in Fig. 4i, the zebrafish cluster dr4 specific *gda* is also highly induced by injury at 1 dpci. Thus, while *cd9b* (*Cd9* in mice) is upregulated in both species, inductions of *gda* and *fn1a* in macrophages are zebrafish specific. Overall, these results indicate both shared and distinct immune responses between regenerative and profibrotic myocardial injury models.

**Fig. 7:**
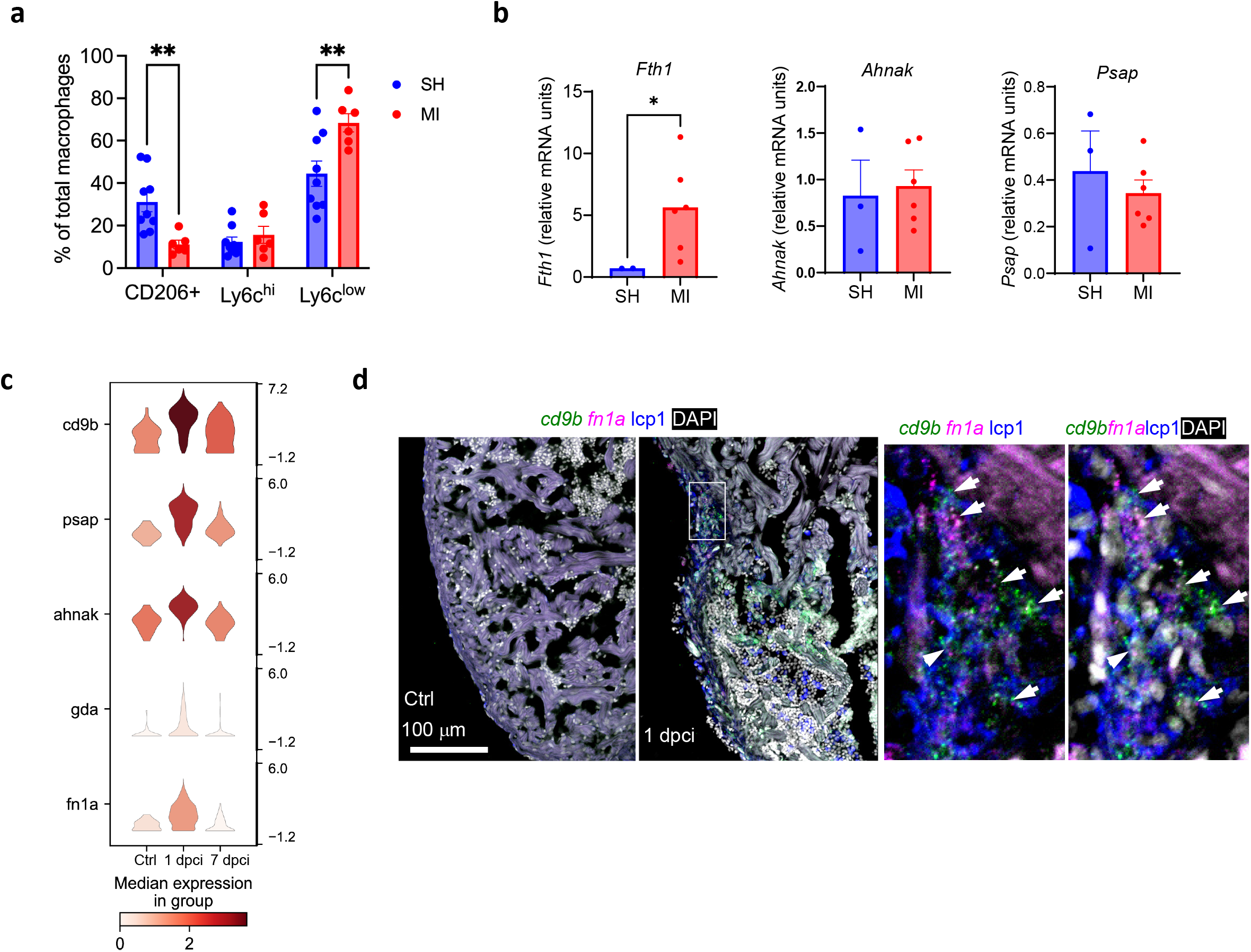
Cardiac macrophage response to heart injury by myeloid clusters in mouse and zebrafish. **a,** Bar plots of the percentage of cardiac macrophages expressing CD206 and Ly6c as determined by flow cytometry in mice undergoing MI or sham at 1 dpi MI. **p<0.01 by Bonferroni’s correction. **b**, Gene expression analyses of Ly6c^low^CD206^-^ FAC-sorted cardiac macrophages from sham or MI groups at 1 dpi. *p<0.05 by Student’s t test. **c**, Violin plots of indicated genes in zebrafish cluster dr4 across timepoints. **d,** Images of heart cryosection showing HCR staining signals of *cd9b* and *fn1a* at 1 dpci in green and magenta, respectively. No expression was detected in the uninjured sample (Ctrl, left panel). Anti-lcp1 staining is shown in blue. DAPI staining is shown in white. The framed region is enlarged to show details on the right with different channel combinations. Arrowheads indicate representative *cd9b*^+^*fn1a*^+^lcp1^+^ cells. Scale bar, 100 μm.

## Discussion

Our study provides the first direct cross-species comparison of immune responses between regenerative and profibrotic myocardial injury models. We also extended our findings and were the first to profile immune cells in multiple extracardiac organs by scRNA-Seq to heart damage in both species. We observed dramatic plasticity in the dynamic immune response that was not limited to the heart. Persistent inflammatory changes, especially in macrophages were noted in extracardiac organs and blood monocytes. Since macrophages are critical for repair after heart injury, this was a major focus of our study. Moreover, we found multiple clusters of macrophages in mice and zebrafish. As expected, there were some unique macrophage subtypes to each species. To our surprise, most of the monocyte/macrophage subclusters from zebrafish aligned to a subcluster in mouse. This permitted us to study analogous pairs of macrophages across species. Our work showed at least at the early 1 dpi time point, the transcriptional response in the analogous macrophage pairs was dramatically different. The vast majority of DEGs were discordant and generally < 10 were concordant. It is thus interesting to hypothesize that disparate stress response may be the key to the fibrotic or healing outcome. At this stage it is unknown if the reaction to stress or the presence of unique regenerative macrophage subclusters in zebrafish or inhibitory macrophage clusters in mouse might be responsible for differences in cardiac healing. How the adult mouse and zebrafish macrophage subclusters compare to neonatal mouse macrophages is another open question. Other groups had previously studied macrophages in non-cardiac tissues after MI. Nahrendorf and colleagues had profiled macrophages in multiple organs mostly by flow cytometry for macrophage subtypes and targeted gene expression^36^. The Dimmeler lab used scRNA-Seq to study the effects of post-MI heart failure on the bone marrow ^37^. Overall, there is abundant evidence that MI and CHF perturb myeloid cell function that can have deleterious effects on many tissues outside of the heart in mammals.

As this project focused on inflammation, cell isolation and scRNA-Seq protocols were adapted for immune cells. There were many hepatocytes, kidney epithelial cells, and pancreatic islet cells sequenced in the mouse and zebrafish. However, there were relatively few cardiomyocytes sequenced due to their size. In general, we found evidence of peripheral organ dysfunction in the liver and kidney but not in the pancreatic islets in mice. Another limitation of our study is that this does not permit robust in silico assessment of ligand-receptor or cell-cell interactions. Although we observed many of the myeloid cell clusters across different time points and organs, we do not formally know if they definitively represent fixed cell subtypes or activation states. It is possible that there is some degree of plasticity observed within a cell cluster or between clusters. Lineage tracing experiments will be needed in the future to answer this. Moreover, further functional validations on the shared and distinct myeloid clusters between species need to be done to demonstrate their contributions to the regenerative capacity. Future experiments to determine how the macrophage subclusters respond to injury at later time points such as 7 and 30 dpi in mice and zebrafish will be of considerable interest. In summary, our results provide new insights into manipulating immunomodulators for cardiac repair.

## MATERIALS AND METHODS

### Mouse Maintenance

Animal procedures were performed according to approved protocols by the Institutional Animal Care and Use Committee (IACUC) at Weill Cornell Medical College. C57BL/6J (Stock #000664) male and female mice were purchased from Jackson laboratories and maintained in plastic cages under a 12/12-h light/dark cycle at constant temperature (22°C) with free access to water and food. Mice were bred in our animal facility.

### LAD ligation procedure

10-12 weeks old male and female C57BL/6J mice were subjected to LAD ligation or sham surgery as described^38^. scRNA-Seq analysis was performed on male mice with other experimental analyses consisting of mice of both sexes. Briefly, mice were anesthetized with isoflurane and then orally intubated and connected to a ventilator for mechanical ventilation. The left pectoralis major muscle was bluntly dissociated until the ribs were exposed. The muscle layers were pulled aside and fixed with an eyelid-retractor. Left thoracotomy was performed between the third and fourth ribs to visualize the anterior surface of the heart and left lung. The pericardium was removed and the proximal segment of the left coronary artery ligated with a 7-0 Ethilon suture. In sham surgeries, the suture step was skipped. The rib cage was closed with stitches on the 3rd and 4th ribs and the skin closed with wound clips. Postoperative analgesia consisted of meloxicam (2 mg/kg) and buprenorphine (0.5 mg/kg). The mice were extubated when spontaneous breathing was observed. The animals were kept in a warm cage until recovery.

### Venipuncture mouse blood collection

For blood cytometry analyses, mouse blood was collected from the tail vein as follows. A small puncture is introduced into the vein with a scalpel. Droplets of blood were collected with a Microvette CB300 (Sarstedt).

### Mouse tissues single cell isolation

Whole blood was collected by cardiac puncture. Red blood cells were lysed with ACK (Ammonium–chloride–potassium) lysis buffer and removed by density centrifugation on a 5% BSA cushion (250g for 5min). Cells were strained with a 40 µm sieve.

For kidney cell isolation, animals were perfused with PBS-5mM EDTA through the left ventricle, then kidneys were collected and minced with a razor blade. Tissue fragments were digested in HBBS – 0.15mg/ml Liberase TM for 25min at 37deg C with vigorous agitation. Digestion was stopped with FBS. The digestion mix was passed through a 70 µm sieve and spun down for 5min at 300g. The pellet was spun down on a 7.5% BSA cushion, for 4min at 250g twice to clear debris.

For heart, non-myocyte cells, hearts were perfused by the Langendorff method as described with minor changes: Collagenase in the digestion buffer was replaced with 0.15mg/ml Liberase TM (Sigma), and only the LV of sham-operated animals or the infarcted area of the LV of infarcted hearts was cut and triturated with a plastic Pasteur pipette. Myocytes were then separated by centrifugation at 50g for 5 mins and discarded. The supernatant with the rest of the cells was pelleted at 300g for 5 mins. Cells were then passed through a 70 µm strainer and debris removed by centrifugating the cells over a 5% BSA-PBS cushion at 250g for 4 minutes twice.

Liver cells were isolated as described^39^. Briefly, mice were anesthetized with ketamine/xylazine (100 mg of ketamine/kg of body weight and 10 mg of xylazine/kg). Livers were perfused *in situ* with liver perfusion medium (Invitrogen) and then digested with liver digestion medium (Invitrogen). Livers were then dissected, placed in ice-cold hepatocyte wash medium (Invitrogen), and the capsule of the liver was then opened to release the cell suspension. Hepatocytes were separated by centrifugation at 30g for 4 minutes. The rest of the cells (supernatant) were pelleted by centrifugating 5 min at 300g. Non-hepatocyte liver cells were passed through a 70 µm cell strainer. Hepatocytes and the rest of the liver cells were combined 1:1.

Pancreatic islets were isolated as described^40^. Briefly, mouse pancreases were perfused with CiZyme (Vitacyte) through the common hepatic duct. Pancreases were removed and digested at 37 °C for 17 min. After two washes with RPMI medium with 3% FBS, islets were separated into a gradient using RPMI medium and Histopaque (Sigma-Aldrich). Islets were then hand-picked and dispersed into a single-cell suspension with a trypsin/EDTA treatment.

### Mouse single cell-RNA sequencing

All cell suspensions were stained with trypan blue and counted using a hemocytometer. Cell suspensions were diluted to target a recovery of 10,000 cells per sample at 50,000 reads per cell. The isolated cells were sent to the Epigenomics Core Facility of Weill Cornell Medicine for single-cell RNA-seq library preparation using the 10x Genomics Chromium Single Cell 3’ GEM, Library & Gel Bead Kit v3, and Chromium Single Cell B Chip Kit. The libraries were sequenced on a pair-end flow cell with a 2 x 50 cycles kit on Illumina NovaSeq6000.

### RNA extraction and qPCR analysis

RNA isolation was performed with Qiagen RNeasy Micro kits. cDNA was synthesized through reverse transcription using a cDNA synthesis kit (Thermo). cDNA was analyzed by real-time PCR using specific gene primers and a SYBR Green Master Mix (Quanta).

### Histology and Immunostaining for Mouse Samples

Freshly collected mouse tissues were fixed with 10% neutral buffered formalin overnight at 4°C. Tissues were then transferred to 70% ethanol and subsequently embedded in paraffin and sectioned at 5 μm thickness. Tissue sections were sent to Weill Cornell’s Laboratory of Comparative Pathology for hematoxylin and eosin (H&E) staining.

For immunohistochemistry (IHC) staining, liver sections were dewaxed, and antigen retrieval was performed using 10 mM sodium citrate buffer (pH 6.0) at boiling temperature for 14 min. Sections were incubated overnight at 4 °C with anti-Mac2 antibody (Cedarlane cat. CL8942AP), followed by incubation with corresponding biotinylated secondary antibodies. HRP-conjugated avidin–biotin complex reagent was used following the manufacturer’s protocol (Vector). Slides were developed using 3,3′-diaminobenzidine (DAB) then counterstained with hematoxylin. Positive cells were then counted manually with an optical microscope.

### Flow Cytometry and Cell Sorting

Single cells were stained with fluorescent antibodies and analyzed and FAC-sorted on a Sony MA900 cell sorter. The following antibodies were used for white blood cell analysis: FITC Anti-CD45R/B220 (Biolegend cat. 103206), APC anti-CD3 (Biolegend cat. 128612), Pacific Blue™ anti-CD4 (Biolegend cat. 100427), PE anti-CD8 (Biolegend cat. 104906), FITC anti-CD11b (Biolegend cat. 101206), Pacific Blue™ anti-Ly-6C (Biolegend cat. 128014), and PE anti-CD11c (Bioleglend cat. 117307). The following antibodies were used for cardiac macrophages: FITC anti-CD11b (Biolegend cat. 101206), PE anti-F4/80 (Biolegend cat. 123110), Pacific Blue™ anti-Ly-6C (Biolegend cat. 128014), and Alexa Fluor® 647 anti-CD206 (MMR) (Biolegend cat. 141712).

### IPA analysis

Ingenuity Pathway Analysis (IPA) version 01-20-04 (Qiagen) was utilized for mouse pathway analyses.

### Zebrafish maintenance

Animal procedures were performed according to approved protocols by the Institutional Animal Care and Use Committee (IACUC) at Weill Cornell Medical College. Adult zebrafish of the Ekkwill strains were maintained as described at 28°C under 14h/10h light/dark cycles^41, 42^. Adult zebrafish (8 month old) of both sexes were used for the experiments. Heart cryosection injury was done as described previously^43^. The *Tg(tcf21:nucEGFP)^pd^*^41^ line^44^ was used. All reporters were analyzed as hemizygotes.

### Zebrafish cell isolation and single cell-RNA sequencing

Two biological replicates were included for all collected samples. Heart ventricles were collected from adult hearts either with cryoinjury or uninjured control fish. Briefly, ventricles were collected as described previously^45, 46^. Ventricles were placed in the ice-cold PBS buffer with 1% BSA. After several washes removing blood cells, ventricles were gently cut into small pieces. Tissues were incubated in digestion buffer (0.5 ml HBSS plus 0.26 U/ml Liberase DH [Roche] and 1% sheep serum) for 45 minutes at 37°C while agitating at 750 rpm on an Eppendorf ThermoMixer. Supernatants were collected every 15 min and neutralized with 10% sheep serum, and cells were completely disaggregated by gently pipetting up and down. The dissociated cells were centrifuged at 200 g for 5 min at 4°C and re-suspended in HBSS with 0.05% BSA. The cell suspension (2 ml) was then gently added on top of 2 ml HBSS buffer with 7.5% BSA. After spinning down at 200 g for 5 min at 4°C, the pellet was resuspended in 500 ml and filtered through a 35 mm strainer. Whole kidney marrow cells were isolated as described previously^47^. Briefly, kidneys were collected on ice and homogenized immediately in PBS plus 0.05% BSA by using a 10ml syringe with 18 1/2 G. The dissociated cells were washed twice in PBS plus 0.05% BSA and spun down at 300 g for 5 min at 4°C. Pellets were re-suspended in PBS plus 0.05% BSA and filtered through a 35 mm strainer. Livers were collected on ice and incubated in digestion buffer (0.5 ml HBSS plus 0.13 U/ml Liberase DH [Roche] and 1% sheep serum) at room temperature. The cell lyses was gently stirred with a Spinbar® magnetic stirring bar (Bel-Art Products), and the supernatant were collected every 5 minutes. The cell lysate was filtered through a 70 mm strainer and centrifuged at 150 g for 3 min at 4°C. Pellet was resuspended in PBS plus 0.05% BSA and then gently added on top of HBSS buffer with 7.5% BSA. After spinning down at 300 g for 5 min, pellets were washed one more time with PBS plus 0.05% BSA. After centrifuging at 300 g for 5 min, the resuspended cell lyses was filtered through a 35 mm strainer. Pancreatic cells were isolated using the same protocol as liver cells except for the initial spinning down at 300 g for 5 min. All suspended cells were stained with trypan blue and counted using a hemocytometer. The isolated cells were sent to the Epigenomics Core Facility of Weill Cornell Medicine for single-cell RNA-seq library preparation using the 10x Genomics Chromium Single Cell 3’ GEM, Library & Gel Bead Kit v3, and Chromium Single Cell B Chip Kit. The libraries were sequenced on a pair-end flow cell with a 2 x 50 cycles kit on Illumina NovaSeq6000. We sequenced 2 replicates of all samples (24 total).

### scRNA-seq analysis

The raw reads were aligned and processed with the CellRanger pipeline (v6.0.0) using the mouse (mm10) or zebrafish genomes (GRCz11). Subsequent analyses were performed in R following the recommendations of Amezquita et al. (https://osca.bioconductor.org/) using numerous functions provided in the R packages scater^48^ and scran^49^. Briefly, quality control was carried out for each sample separately with functions from the scuttle^48^ package; cells with low gene content and high mitochondrial gene content were removed from further analyses. For each species, different count matrices across all samples were then scaled for sequencing depth differences and log-transformed using multiBatchNorm from the batchelor ^50^ package. Cell types were annotated with SingleR^25^ using numerous datasets and manual inspection of marker genes. Data was also explored and annotated with Cellxgene version v0.16.8. scRNA-seq data UMAPs, SAMaps, dot plots, and violin plots were generated with Cellxgene VIP (https://doi.org/10.1101/2020.08.28.270652).

### Cross-species comparisons

SAMap^35^ (v1.0.2) was used to determine cell homology between mouse and zebrafish myeloid clusters. Briefly, a reciprocal BLAST map between mouse and zebrafish transcriptomes was generated using the map_genes.sh script from SAMap, which ran tblastx (nucleotide-nucleotide BLAST, NCBI) in both directions with respect to Ensembl zebrafish and mouse transcriptomes (Danio_rerio.GRCz11.cds.all.fa and Mus_musculus.GRCm38.cds.all.fa). Raw scRNA-seq count matrices of zebrafish and mouse were extracted from SingleCellExperiment objects with cluster annotations and converted to h5ad format using the sceasy R package (v0.0.7). Raw data were then processed and integrated by SAMap. Mapping scores between the myeloid clusters of each species were calculated with SAMap get_mapping_scores (n top=0) and visualized in R using ggalluvial; cluster pairs with alignment scores less than 0.2 were excluded.

For myeloid cluster pairs with high inter-species alignment scores (mm1-dr4, mm4-dr6, and mm6-dr3), marker genes for each cluster were assessed using scran::scoreMarkers (mean AUC > 0.6), and over-representation analyses were performed using clusterProfiler^51^ with respect to gene ontology terms. To compare the transcriptional response to MI (mouse) or cardiac cryoinjury (zebrafish) between the myeloid cluster pairs, differential expression analysis was performed using scran::findMarkers(lfc=0.25) at days 1 and 7 in the heart and in the blood. Genes were considered statistically significant at an FDR cutoff of 0.10.

### Histology and Microscopy

Freshly collected zebrafish tissues were fixed with 4% paraformaldehyde (PFA) overnight at 4C and applied to cryosection at a 10 mm thickness. Hybridization Chain Reaction (HCR 3.0) staining of whole-mounted hearts or cryosections was done following the published protocols^33^. HCR probes for *havcr1*, *moxd1*, *cd9b*, *fn1a*, and *gda* were synthesized by Molecular Instruments Inc. For immunostaining of whole-mounted hearts or heart sections^52, 53^, samples were blocked with 2% bovine serum albumin (BSA, VWR, cat#97061), 1% DMSO, 0.5% goat serum (ThermoFisher, cat#16210) and 0.5% Triton X-100 in PBS for 1 hr at room temperature (RT). Primary antibodies were diluted in the blocking buffer and incubated with hearts overnight at 4C. Hearts were then washed with PBS plus 0.1% Tween 20 and incubated with the secondary antibody diluted in the blocking buffer for 1.5 hr at room temperature. Hearts were stained with DAPI (ThermoFisher, D3571) to visualize nuclei. Primary antibodies used in this study were rabbit anti-lcp1 (GeneTex GTX124420, 1:200), Secondary antibody used in this study were Alexa Fluor 488 goat anti-rabbit (ThermoFisher, 1:200). All antibodies are commercially available and are validated by suppliers. Fluorescent images of tissue sections were captured used a Zeiss 800 confocal microscope (Zen 2.6 blue edition software).

### Statistics and Reproducibility

Unless otherwise stated, data are presented as mean ± s.e.m. Data are derived from multiple experiments unless stated otherwise. If not mentioned otherwise in the figure legend, statistical significance is indicated by \**P* < 0.05, \*\**P* < 0.01 and \*\*\**P* < 0.001. Statistical analysis was carried out using unpaired, two-tailed *t*-test, one-way, or two-way ANOVA with Tukey’s honest significant difference (HSD) post hoc test. GraphPad Prism 7 was used for statistical analysis.

## DATA AVAILABILITY

The scRNA-seq datasets generated in this study have been deposited at NCBI’s Gene Expression Omnibus under accession number GSE227191. This will be made publicly available upon journal submission.

## CODE AVAILABILITY

All scripts as well as the code and cell labels used for generating the scRNA-seq based figures will be made available upon journal submission.

## ACKNOWLEDGMENTS

We thank Adedeji A. Afolalu, Chaim Shapiro, Soji Hosten, and Chelsea Quaies for fish care. This work was supported by Rudin Foundation fellowships to Y.X. and J.Y., a predoctoral training grant position in Stem Cell Biology and Regenerative Medicine from New York State Stem Cell Science program (NYSTEM) to B.P. The research was supported by Chan Zuckerberg Initiative grant (DAF2020-217734) to D.B., J.C, and J.C.L., Weill Cornell Start-up funds and NIH grant (R01HL155607) to J.C.

## AUTHOR CONTRIBUTIONS

E.C., J.Y., D.B., J.C., and J.C.L. designed the study and wrote the manuscript with input from all authors. E.C., J.Y., Y.X., A.R.-N., B.Y., B.P., M.Q., E.A.H., L.S., performed and analyzed the animal experiments. P.Z., F.D., and A.M.P. analyzed the scRNA-Seq experiments. J.C.L, J.C., and D.B. conceived and supervised the study and acquired funding for the work.

## COMPETING INTERESTS

The authors declare no competing interests.

## Figure Legends

**Extended Data Figure 1.**
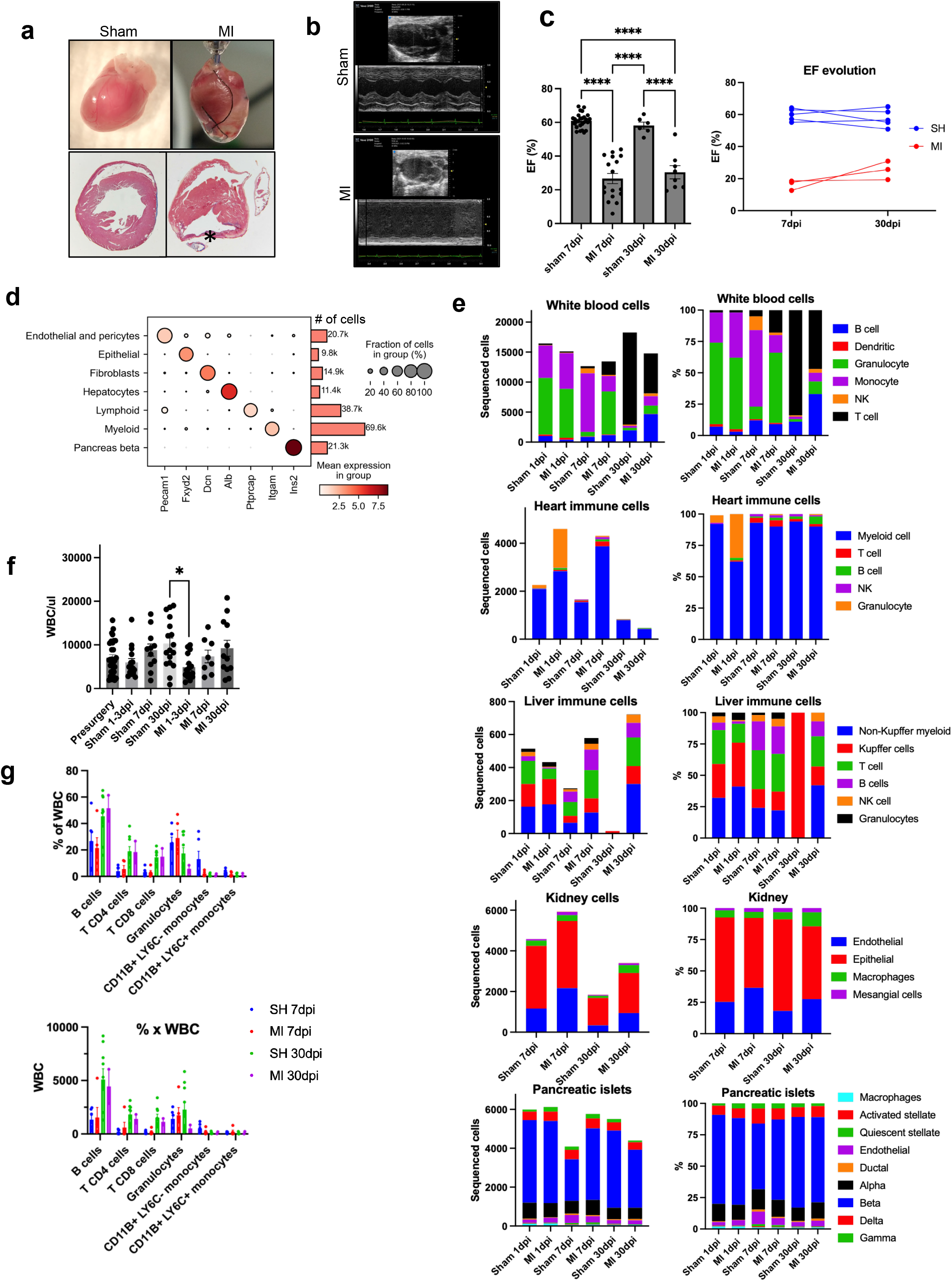
Multiorgan cellular response to MI. **a,** Top: Representative pictures of hearts 30 dpi after excision. Bottom: Representative Masson’s Trichrome staining of formalin fixed paraffin embedded 30 dpi heart sections. Asterisk marks the infarction site. MI: myocardial infarction; SH: sham. **b,** Representative Motion-mode Parasternal Long Axis echocardiography frames at 30 dpi. In the MI (bottom), there is akinesis of the anterior left ventricle (LV) wall. **c,** Quantitation of the left ventricular ejection fraction (LVEF) of sham and MI at 7 and 30 dpi (Sham 7 dpi *n*=26; MI 7 dpi *n*=17; Sham 30 dpi *n*=6; MI 30 dpi *n*=8) (left). LVEF evolution in individual mice analyzed at 7 and 30 dpi. **d,** Dot plot of major cell types, regardless of tissue, time point or surgery type, and their top marker genes. Color intensity represents normalized mean expression. Size of the dot represents fraction of cells with expression of the given gene. **e,** Bar plots on the left show number of sequenced cells included in the study, by sample and tissue. Bars are colored by cell type composition. Bar plots on the right show the percentage of cells in each sample and tissue. **f,** WBC counts per ul of blood of mice (Presurgery *n*=30; Sham 1-3 dpi *n*=15; Sham 7 dpi *n*=11; Sham 30 dpi *n*= 17; MI 1-3 dpi *n*=16; MI 7 dpi *n*=8; MI 30 dpi *n*=12). **g,** Frequencies of WBC (top) and absolute counts (bottom) of WBC by flow cytometry of mice undergoing sham or MI procedure at 7 and 30 dpi (Sham 7 dpi *n*=6; MI 7 dpi *n*=5; Sham 30 dpi *n*=9; MI 30 dpi *n*=3). **e-g**, Data are mean +/-SEM. A one-way ANOVA followed by Tukey’s Honest Significant Difference post hoc test was used for statistical analysis (\**p*<0.05; \*\*\*\**p*<0.0001).

**Extended Data Figure 2.**
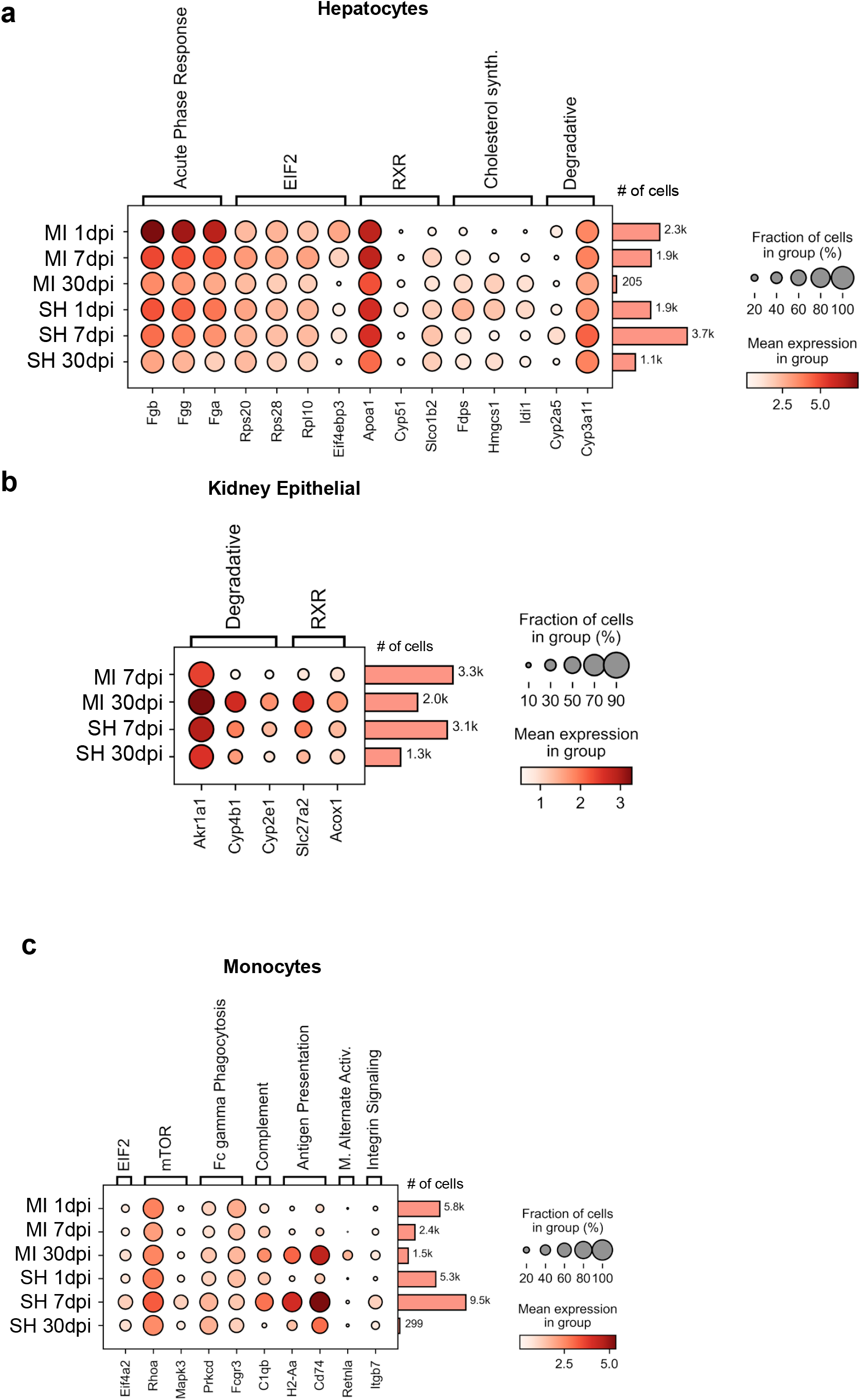
Ingenuity pathway target gene expression. Dot plot shows gene expression in hepatocytes (**a**), heart endothelial cells (**b**), and kidney epithelial cells (**c**) by experimental group (MI or sham) and time point. Color intensity represents normalized mean expression. MI: myocardial infarction; SH: sham.

**Extended Data Figure 3.**
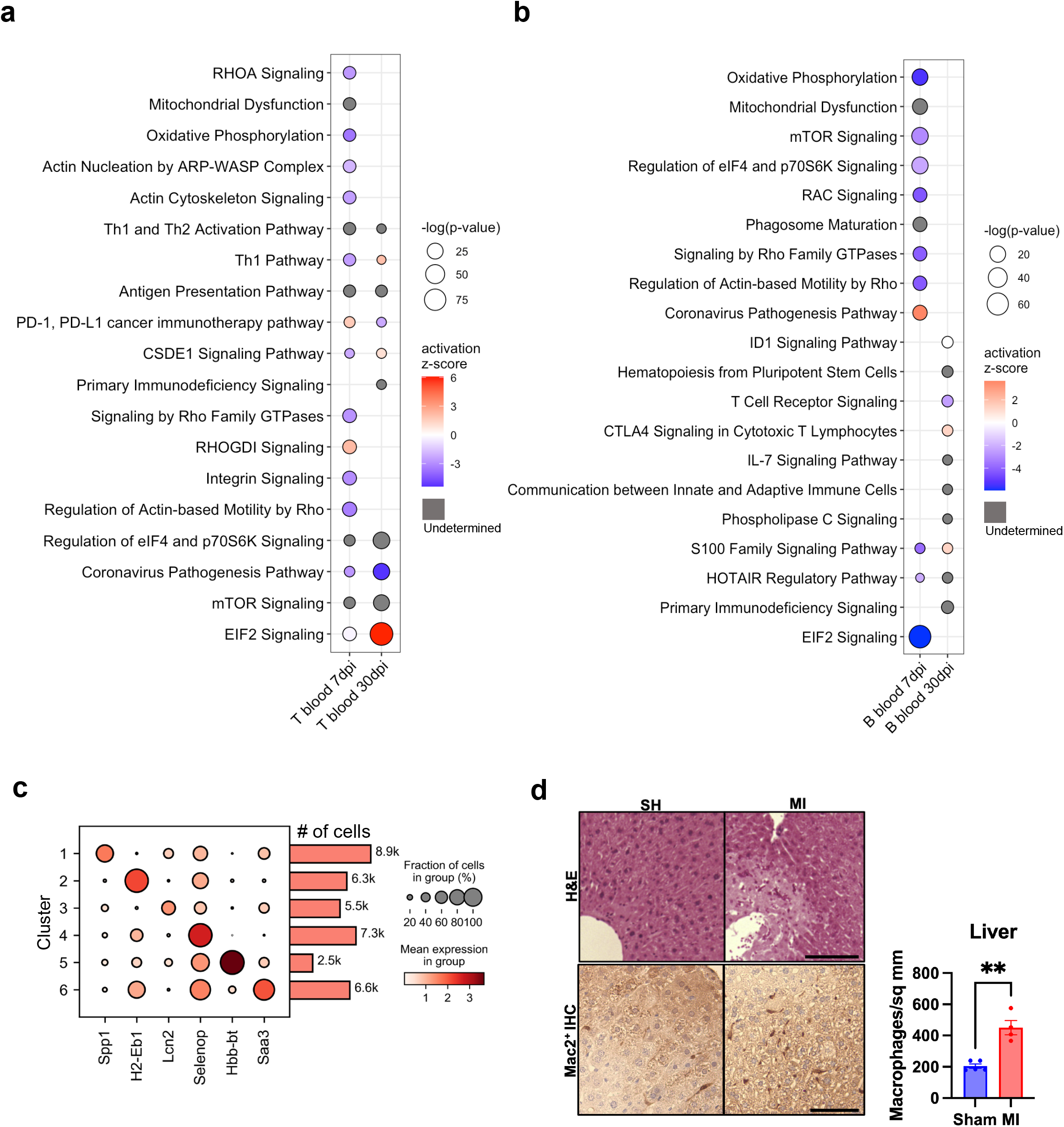
Immune cell response to MI in blood and liver by scRNA-Seq. **a,b** Dot plots depicting enriched pathways in MI compared to sham and their statistical significance and activation z-score of blood T (**a**) and B (**b**) cells at 7 and 30 dpi. Positive z-score predicts activation of the pathway in the MI compared to the sham group. Negative z-score predicts inhibition of the pathway in the MI compared to the sham group. Gray dots denote undetermined activation status. **c,** Dot plot of the top marker genes for the myeloid clusters. Color intensity in dot plots represents normalized mean expression. **d,** Representative liver sections from mice in the sham and MI group at 30 dpi. H&E staining (top, scale bar is 0.5 mm) and Mac-2 immunohistochemistry staining (bottom, scale bar is 50 µm) for macrophages. Right panel: Quantification of Mac2+ liver macrophages. Data are presented as mean +/-SEM (Sham *n*=5; MI *n*=4 male mice). A two-tailed Student’s t test was used for statistical analysis (\*\**p*<0.01).

**Extended Data Figure 4:**
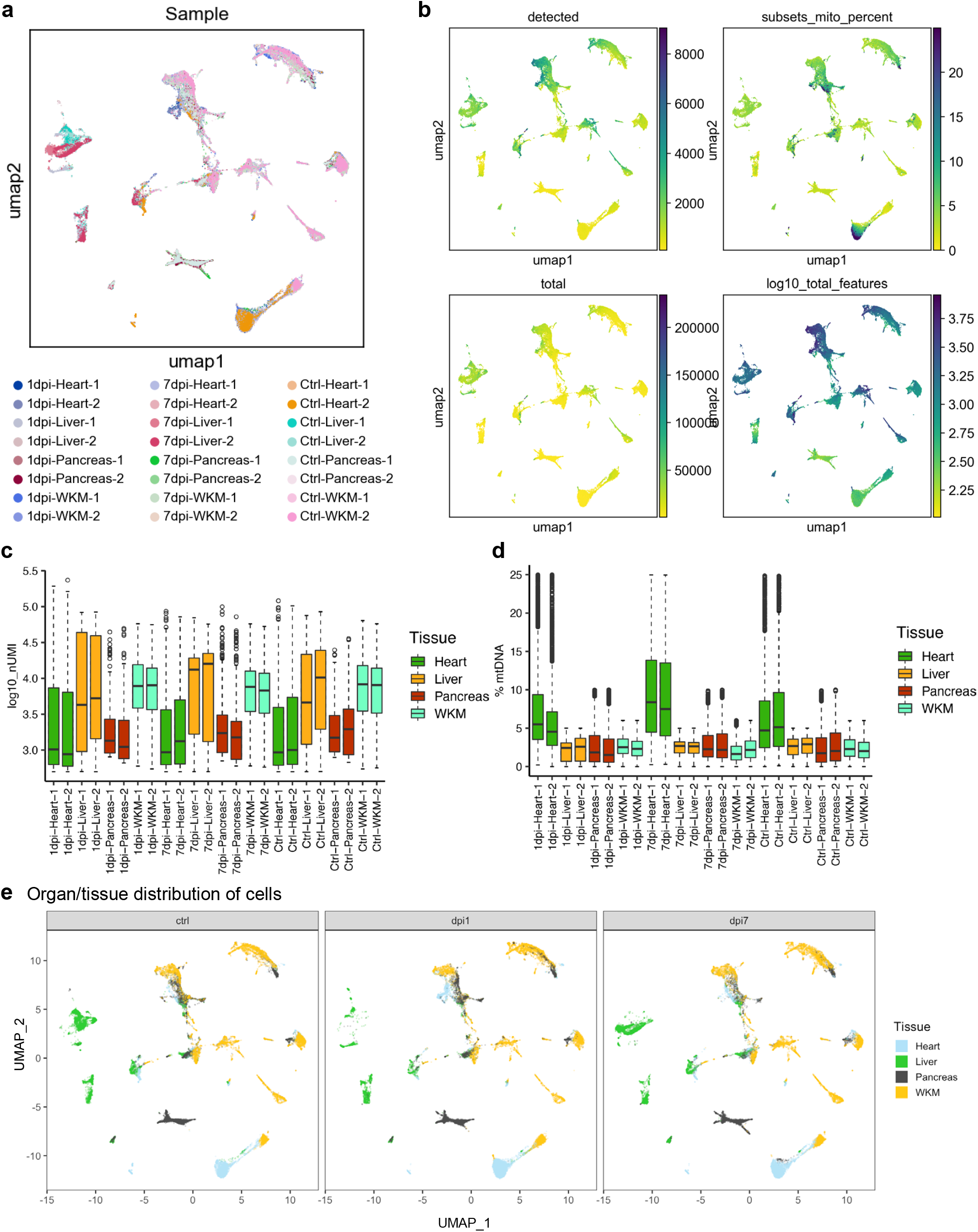
Sample distribution and QC of zebrafish scRNA-seq. **a,** A UMAP showing the distribution of 24 samples over clusters. **b,** UMAPs depicting the number of detected features (detected), percentage of mitochondria transcripts (subset_mito_percent), the total number of transcripts (total), and log10 converted total number of transcripts (log10_total_features). **c,** Box plots showing the number of detected features (left) and percentage of mitochondria transcripts (right) across 24 samples. **d,** UMAPs showing tissue membership of clusters across treatment groups (Ctrl, 1 dpci, and 7 dpci).

**Extended Data Figure 5:**
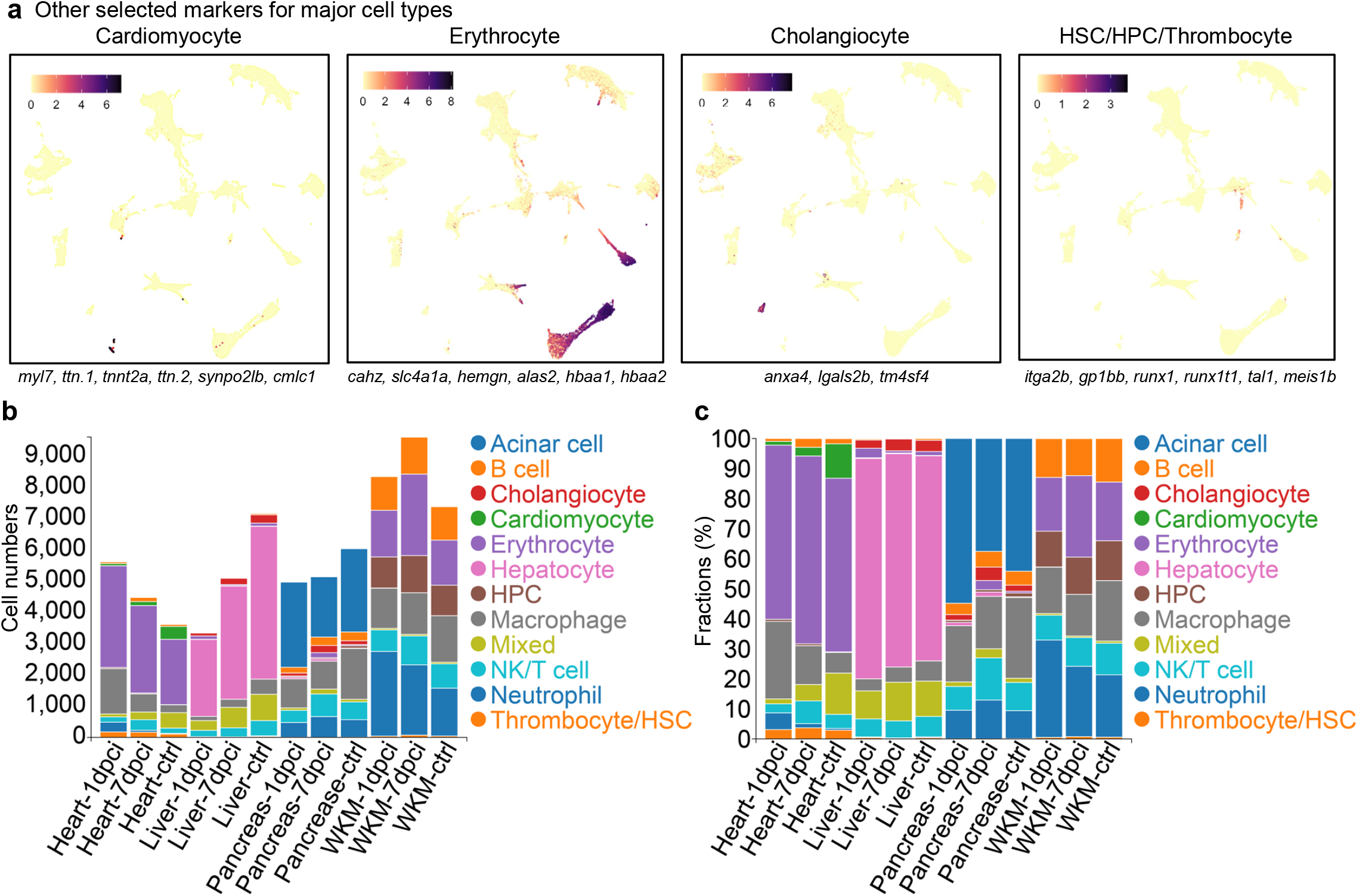
Major cell types detected in the zebrafish single-cell dataset. **a,** UMAPs demonstrating expressions of groups of marker genes for cardiomyocyte, erythrocyte, cholangiocyte, and HSC/HPC/Thrombocyte, respectively. **b,c,** Absolute cell count (b) and cell fraction (c) of major cell types across 24 samples.

**Extended Data Figure 6:**
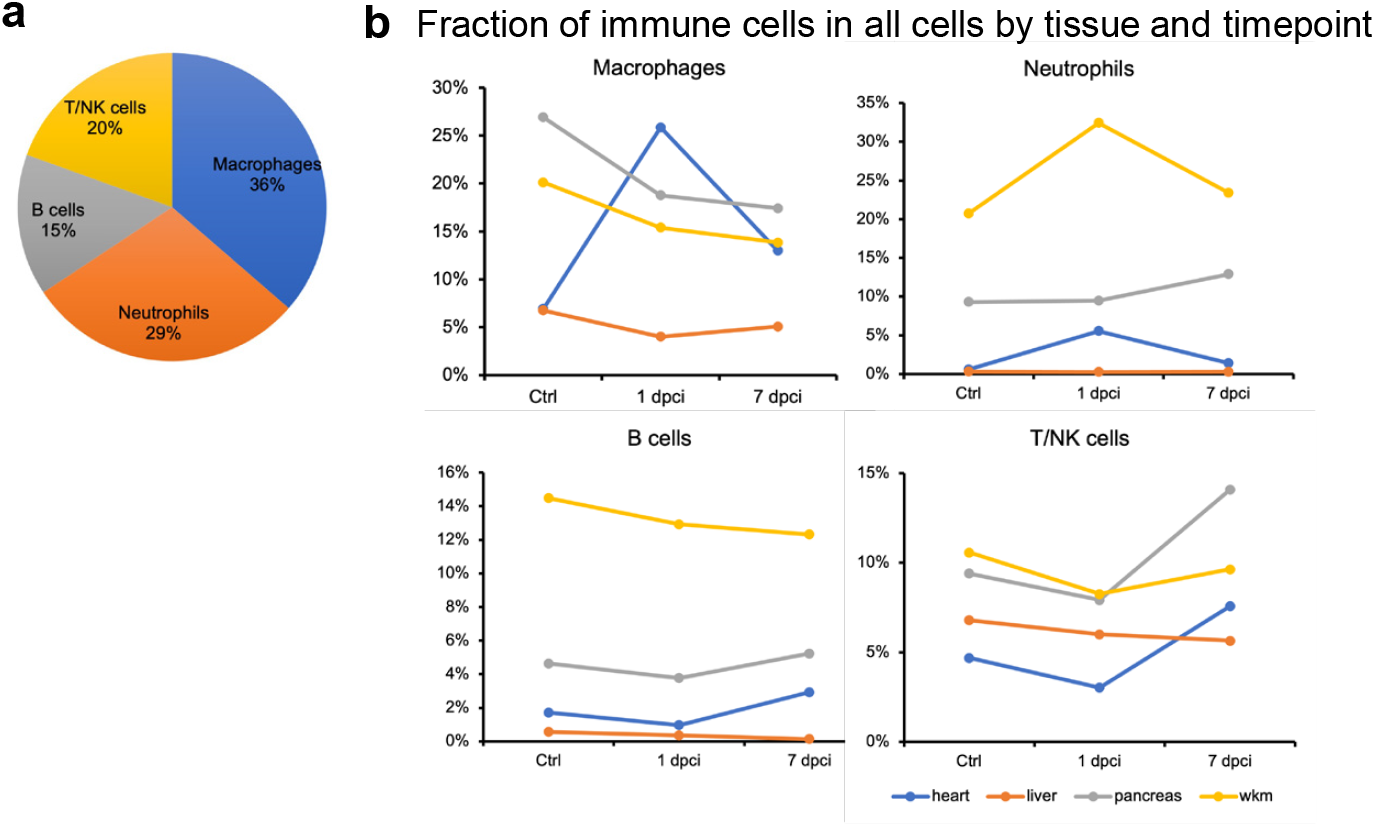
Characterization of major immune cells in the zebrafish single-cell dataset. **a,** Fractions of major immune cells in all samples. **b,** Dynamics of immune cell fractions over all cells in each tissue and treatment group (Ctrl, 1 dpci, or 7 dpci).

**Extended Data Figure 7:**
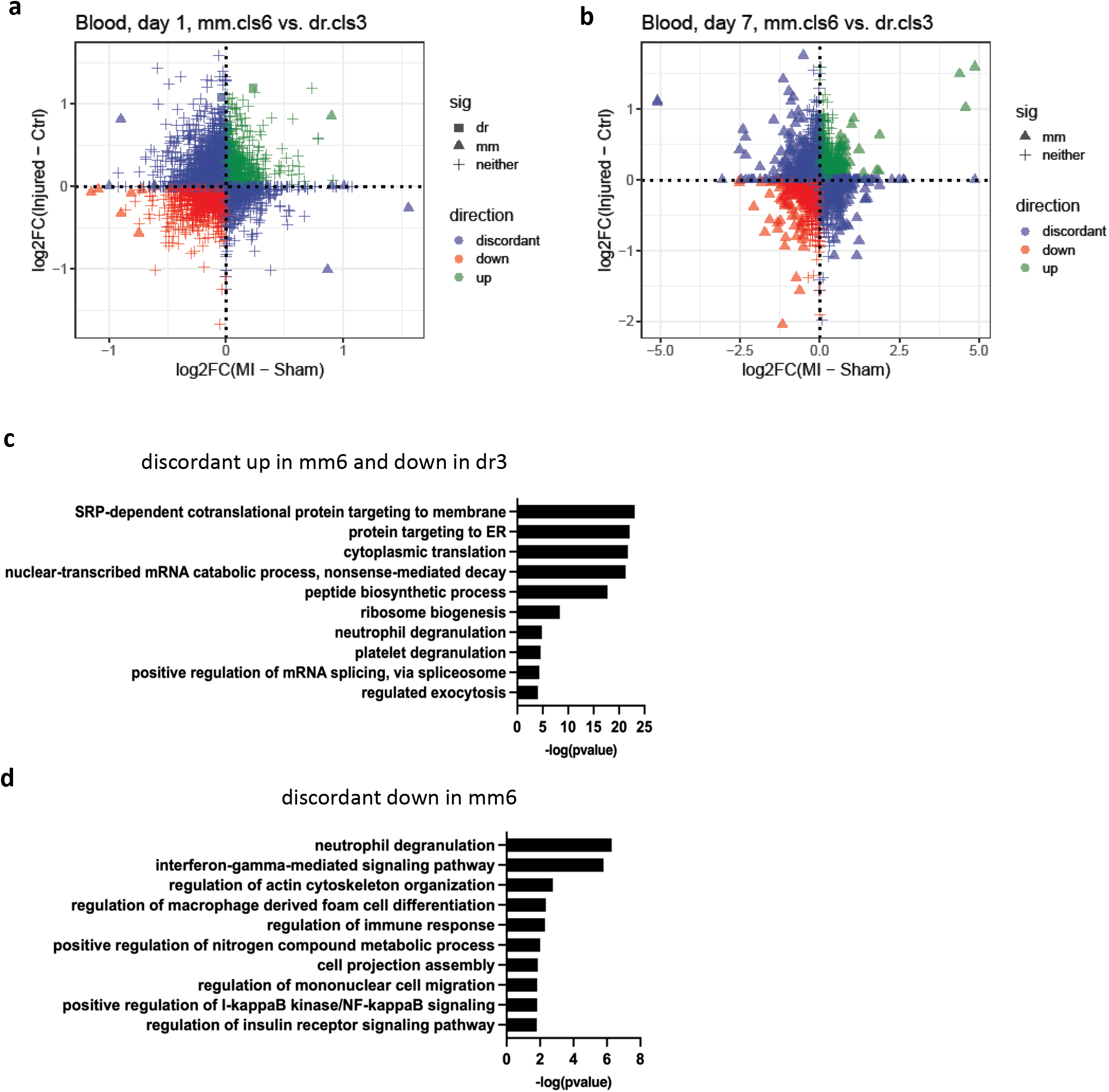
Comparison of the analogous murine and zebrafish myeloid clusters. **a, b,** Scatterplot of the fold changes in murine and zebrafish genes with cardiac injury compared to control in mm6 and dr3 blood monocytes cells at 1 (**a**) and 7 (**b**) dpi (FDR<0.10 and |log2FC|>0.25). **c, d,** Enriched biological process terms of the DEGs that are discordant between mm6 and dr4 cells in **a**.

**Extended Data Figure 8:**
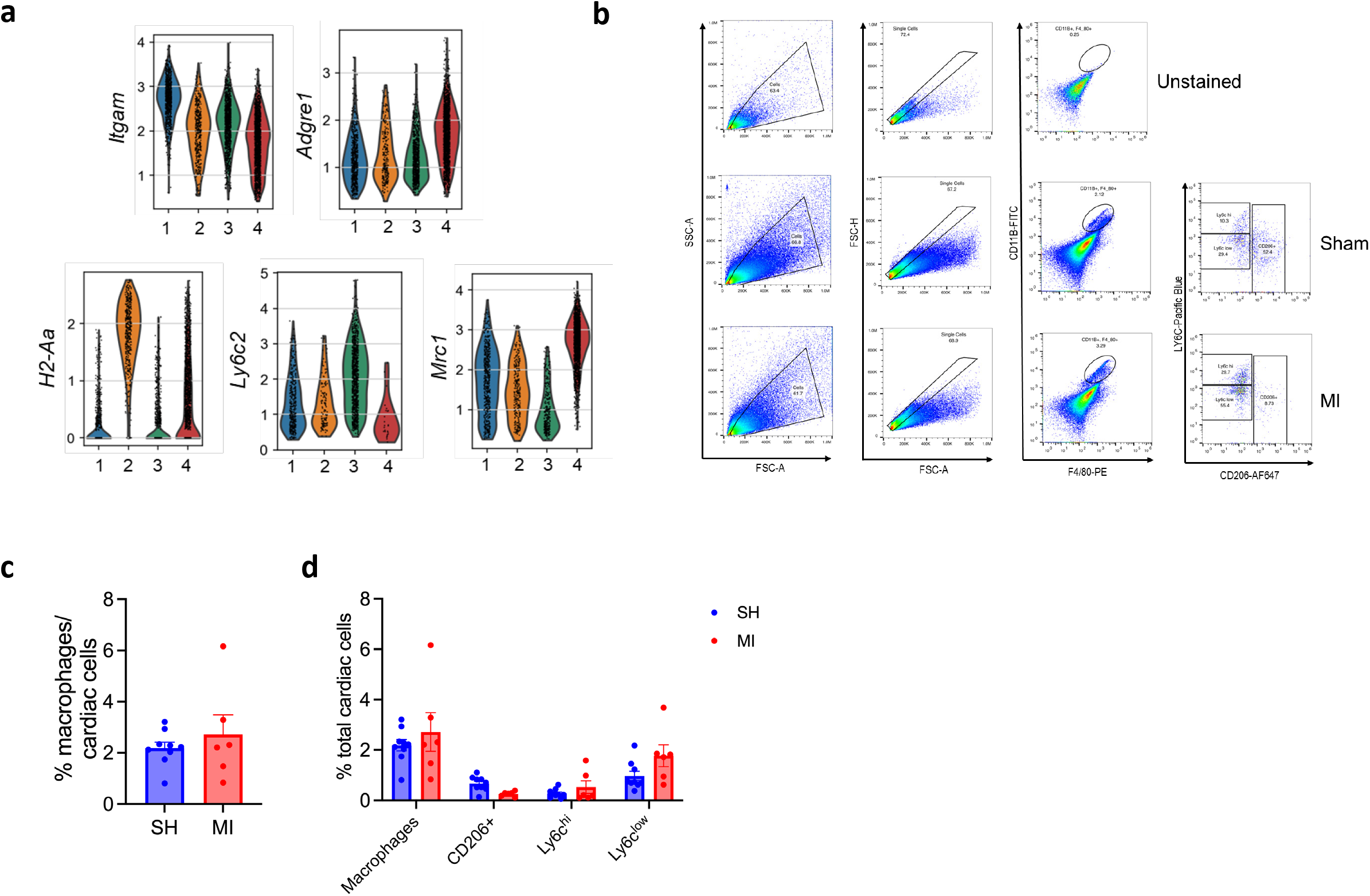
Changes in myeloid subclusters in mouse and zebrafish with cardiac injury. **a,** Violin plots of indicated myeloid genes according to cardiac myeloid subclusters in mouse at 1 dpi. **b,** Representative flow cytometry gating strategy for cardiac macrophages in mice undergoing sham or MI at day 1 (FDR<0.10 and |log2FC|>0.25). **c-d,** Enumeration of cardiac macrophages in mice with sham (SH) or myocardial infarction (MI) procedure at day 1 plotted as percentage of all heart cells (**c**) or total numbers (**d**).

## Notes

### Competing Interest Statement

The authors have declared no competing interest.

